# Magnesium modulates *Bacillus subtilis* cell division frequency

**DOI:** 10.1101/2022.10.01.510448

**Authors:** Tingfeng Guo, Jennifer K. Herman

## Abstract

By chance, we discovered a window of extracellular magnesium (Mg^2+^) availability that modulates *Bacillus subtilis* division frequency without affecting growth rate. In this window, cells grown with excess Mg^2+^ produce shorter cells than those grown in unsupplemented medium. The Mg^2+^-responsive adjustment in cell length occurs in both rich and minimal media and in domesticated and undomesticated strains. Of other divalent cations tested, manganese (Mn^2+^) and zinc (Zn^2+^) also resulted in cell shortening, but only at concentrations that affected growth. Cell length decreased proportionally with increasing Mg^2+^ from 0.2 mM to 2.0 mM, with little or no detectable change in labile, intracellular Mg^2+^ based on a riboswitch reporter. Cells grown in excess Mg^2+^ had fewer nucleoids and possessed more FtsZ-rings per unit cell length, consistent with increased division frequency. Remarkably, when shifting cells from unsupplemented to supplemented medium, more than half of the cell length decrease occurred in the first 10 min, consistent with rapid division onset. Relative to unsupplemented cells, cells growing at steady-state with excess Mg^2+^ showed enhanced expression of a large number of SigB-regulated genes and activation of the Fur, MntR, and Zur regulons. Thus, by manipulating the availability of one nutrient, we were able to uncouple growth rate from division frequency and identify transcriptional changes suggesting cell division is accompanied by oxidative stress and an enhanced demand to sequester and/or increase uptake of iron, Mn^2+^, and Zn^2+^.

**IMPORTANCE:** The signals cells use to trigger cell division are unknown. Although division is often considered intrinsic to the cell-cycle, microorganisms can continue to grow and repeat rounds of DNA replication without dividing, indicating cycles of division can be skipped. Here we show that by manipulating a single nutrient, Mg^2+^, cell division can be uncoupled from growth rate. This finding can be applied to investigate the nature of the cell division signal(s).

## INTRODUCTION

The abundant cation Mg^2+^ is perhaps most appreciated for its role as an enzymatic co-factor, supporting catalysis in hundreds of biochemical reactions (1-3). However, Mg^2+^ has diverse biological functions and is also critical for ribosome assembly (4, 5), chelation and stabilization of ATP and other polyphosphates (6), phosphate uptake regulation (7, 8), osmotic adaptation (9), setting circadian period in plants (10), and supporting envelope integrity in bacteria (11-13). Due to Mg^2+^’s central place in physiology, cells must be able to respond rapidly when availability fluctuates - sometimes over several orders of magnitude, depending on environment. In human serum, where Mg^2+^ concentration is tightly controlled, 0.7 to 1.0 mM is considered homeostatic (14). In contrast, Mg^2+^ in the digestive tract of animals, or in soil and aquatic environments is much more variable, ranging from micromolar to millimolar levels.

While free-living bacteria are able to adapt to large fluctuations in extracellular Mg^2+^, they keep intracellular levels relatively constant. Under replete conditions, cell associated Mg^2+^ in *E. coli* is estimated to be 20-100 mM (15, 16), of which only 1-10 mM is considered labile (16-19). Of the remaining pool, approximately half is associated with nucleic acid, proteins, and ribosomes. The other half is found complexed with the enzymatically relevant form of ATP, the relatively stable chelate Mg^2+^-ATP (20). Not surprisingly, ATP synthesis is tightly coordinated with Mg^2+^ availability (6). In fact, cells will scavenge Mg^2+^ from ribosomes at the expense of protein synthesis before allowing intracellular Mg^2+^ to fall to levels insufficient to support ATP chelation (6, 9, 21).

Intracellular Mg^2+^ is acquired using importer proteins that, in Gram-negatives, are often regulated by a PhoPQ two-component systems (22, 23). These systems sense and respond to changes in both external Mg^2+^ availability and cellular demand in large part by regulating transcription of Mg^2+^ transporters. *B. subtilis* lacks a homologous two-component system and controls expression of its major Mg^2+^ importer (MgtE) using a riboswitch. The M-box riboswitch attenuates transcription of *mgtE* when intracellular Mg^2+^ is sufficient (4). Two other importers, YfjQ and CitM also contribute to Mg^2+^ uptake (24). YfjQ is a minor importer, while CitM is a symporter that allows co-transport of Mg^2+^ and citrate (25, 26). Under hyperosmotic conditions, and concomitant with potassium influx, *B. subtilis* can also efflux Mg^2+^ through the an exporter called MpfA (27).

Aside from its role in supporting basic physiological functions, Mg^2+^ is known to suppress phenotypes associated with inactivation of cell envelope-related genes. Providing 10-25 mM Mg^2+^ (more than a magnitude higher than concentrations found in typical media), restores both viability and rod shape to strains with deletions in the morphogenes *mreB, mbl, mreBH, mreC, and mreD* (28, 29). Mg^2+^ restores rod shape to strains with deletions in *ponA* (encoding PBP1A) and *lytE* (a major D,L-endopeptidase)(11, 30), and to mutants with deletions in the teichoic acid synthesis genes *pgcA, gtaB*, and *ugtP* (31, 32). A Δ*glmR* mutant, which is unable to upregulate gluconeogenesis, is inviable when grown on a gluconeogenic carbon source. Remarkably, millimolar Mg^2+^ can suppress this lethality (4). Mg^2+^ even increases resistance to the cell-wall targeting antibiotic methicillin (33). Thus, Mg^2+^ elicits cellular changes that allow it to function as general suppressor of wide variety of envelope-related defects.

The mechanism by which Mg^2+^ promotes envelope integrity is unclear. *B. subtilis* grown with higher levels of Mg^2+^ have lower levels of amidated meso-diaminopimelic acid in their peptidoglycan (PG)(34); however, reduced amidation is unlikely to account for Mg^2+^ rescue, as a mutant lacking the modification (Δ*asnB*) itself has shape defects that are suppressed by Mg^2+^ (34). Mg^2+^ also reduces the dysregulated D,L-endopeptidase activity associated with deletion of *mreB* (35); it is unknown if Mg^2+^’s impact on endopeptidase activity is direct or a consequence of other effects Mg^2+^ has on the cell.

Here we investigate a phenotype not previously associated with Mg^2+^. We identify a window where increasing Mg^2+^ availability increases cell division frequency without affecting growth rate. Our results suggest that as Mg^2+^ availability decreases, cells prioritize maintenance of cell elongation and linear growth over cell division. This reprioritization of cell resources, which transcriptional profiling suggests is accompanied by changes in metal homeostasis, results in longer cells with more nucleoids and fewer Z-rings.

## RESULTS

### Mg^2+^ supplementation leads to cell shortening in rich media in both domestic and undomesticated B. subtilis strains

In 2017 we obtained a new bottle of premade LB-Lennox powder from Sigma. While examining micrographs of membrane-stained *B. subtilis* 168 cells grown in liquid LB made from the new powder (LB*), we noted the cells appeared qualitatively longer compared to cells grown in previous lots of LB. LB is a rich medium consisting of only NaCl, tryptone, and yeast extract, so we reasoned the longer cells most likely resulted from a difference in trace element content. Mg^2+^ was a priority candidate both because tryptone-based medias like LB tend to be low in Mg^2+^ (35, 36) and because prior studies had shown low Mg^2+^ media results in *B. subtilis* filamentation (37-39); direct comparison to the previously observed filamentation phenotypes was not possible as the microscopy methods did not allow for visualization of septa present along filaments. To test if the longer cells could be the result of low Mg^2+^ content in the new LB, we supplemented the medium with 10.0 mM MgCl^2^ and imaged cells following membrane staining. As shown in Fig 1A and 1B, cells grown with supplemental Mg^2+^ were 36% shorter on average, consistent with the possibility that cells were longer in the new medium due to reduced Mg^2+^ content.

**Figure 1.**
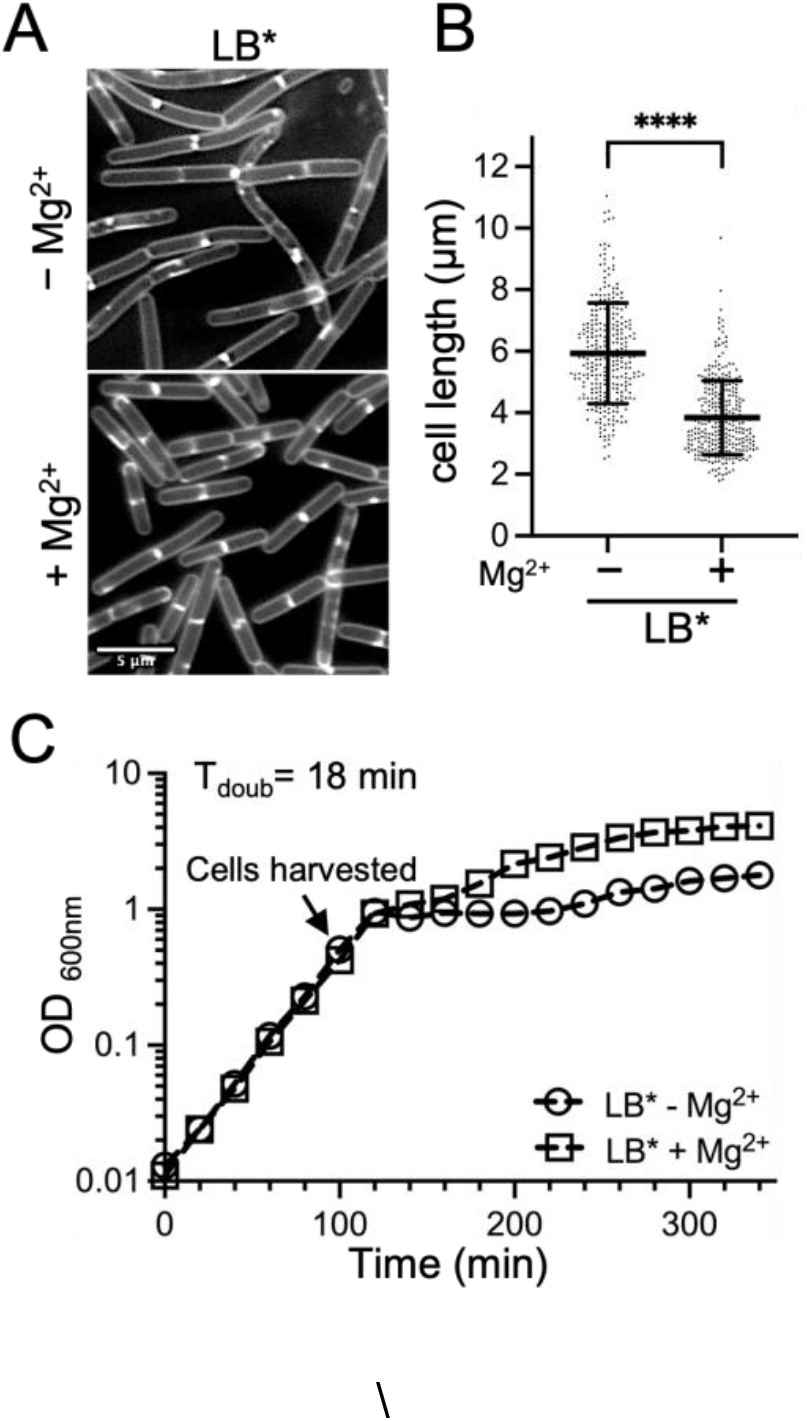
Cell length following growth in LB. WT *B. subtilis* 168 (BJH004) was grown at 37°C to mid-exponential in LB* medium without or with 10.0 mM MgCl_2._ supplementation. The (*) indicates the phenotype was lot-specific and not generalizable to all LB media. (A) Micrographs following membrane staining and epifluorescent microscopy (scaled identically). (B) Scatter plots showing the distribution of cell lengths quantitated for 300 cells from each condition. Bars represent the means of 300 cells ± SD (****, p<=0.0001). (C) Representative growth curves.

Since growth rate can affect cell size (40-46), we next assessed if the Mg^2+^ addition affected the doubling time of cells in the new LB. We found the growth rates of cells cultured without and with 10.0 mM MgCl^2^ were identical during exponential stage (Fig 1C), the same phase of growth the cells were imaged in. From these results, we conclude that there is a growth-rate independent effect of Mg^2+^ availability on cell length, at least in the window tested.

The phenotype we initially observed was specific to only one batch of LB, so next we investigated if the Mg^2+^ responsive phenotype could be observed in another rich medium. CH is a standard complex medium used in *B. subtilis* studies. The casein hydrolysate base (CH*) was sufficient to support robust growth without the standard supplementation of Mg^2+^ and Mn^2+^ salts (Fig 2A), allowing us to utilize CH* for the experiments. Similar to the results with LB*, cells grown in CH* exhibited a Mg^2+^-responsive phenotype. *B. subtilis* 168 (168) cells cultured in CH* with supplemental Mg^2+^ were approximately 2-fold reduced in average cell length compared to CH* only (Fig 2B and 2C). No difference in doubling time (Fig 2A) or cell width (Fig 2D) was detected between CH* and CH* supplemented with 10.0 mM MgCl^2^.

**Figure 2.**
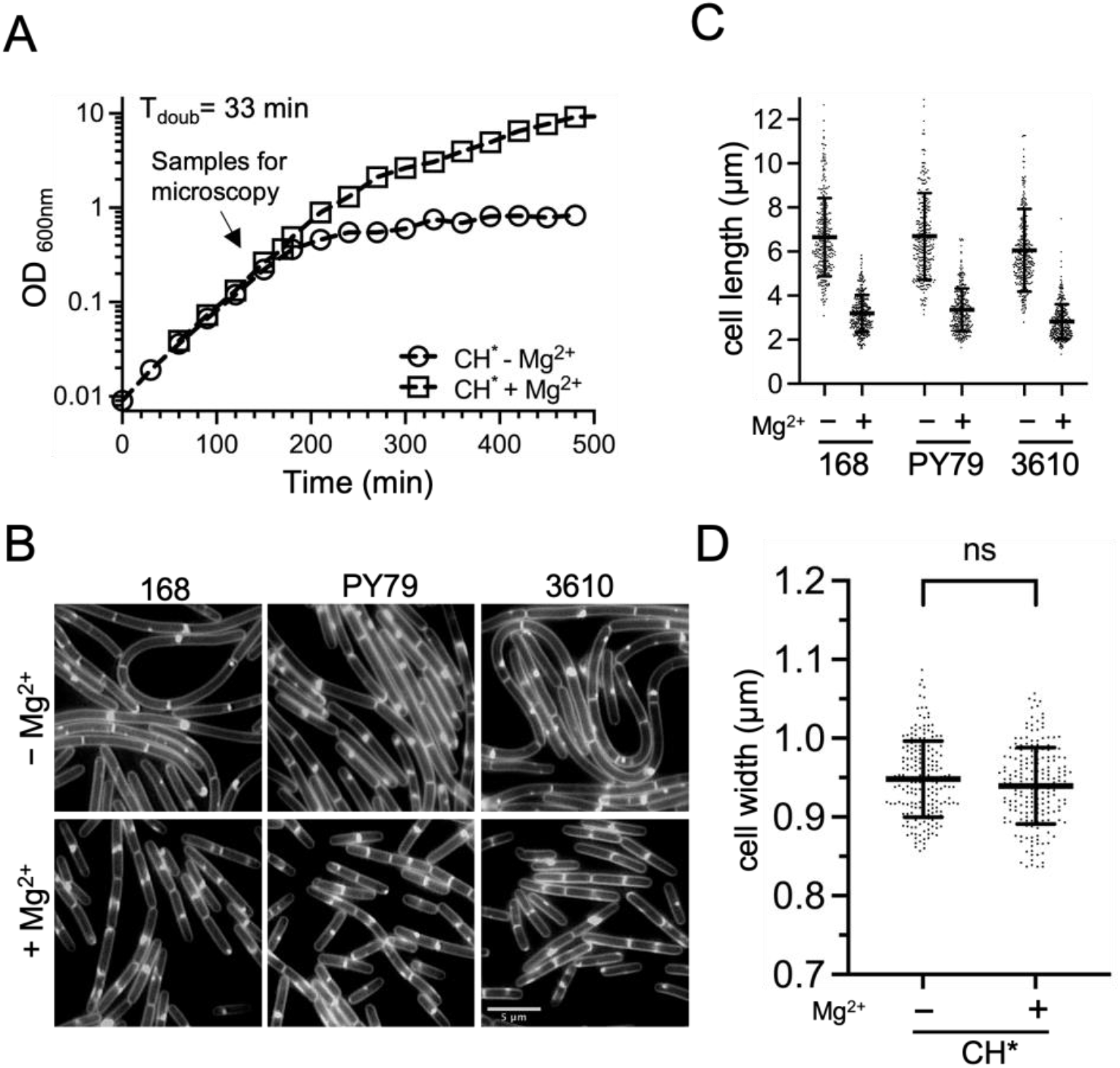
Cell length of three *B. subtilis* strains following growth in CH* medium. WT *B. subtilis* 168 (BJH004), PY79 (BJH001) and 3610 (BJH405) were grown at 37°C to mid-exponential in CH* supplemented with 10.0 mM MgCl_2_ when indicated. (A) Representative growth curves for WT. (B) Representative micrographs following membrane staining and epifluorescent microscopy (scaled identically). (C) Scatter plots showing the distribution of cell with quantitated for 300 cells from each condition. Bars represent the means of 300 cells ± SD. (D) Scatter plots showing the distribution of cell widths for 200 cells grown without or with 10.0 mM MgCl_2_. Bars represent the means of 200 cells ± SD (ns, p>0.05).

We next investigated if the Mg^2+^-responsive phenotype observed in 168 was present in two other commonly utilized *B. subtilis* strains: the prophage-cured laboratory strain, PY79 and the undomesticated wild-type strain, 3610 (47). Independent of strain, the addition of 10.0 mM MgCl^2^ to CH* consistently resulted in cells that were ∼2-fold reduced in length compared to CH* alone (Fig 2B and 2C). These results suggest the effect of Mg^2+^ is likely to be a phenomenon generalizable to the species.

### The Mg^2+^-responsive phenotype occurs in minimal medium and does not require the addition of exogenous amino acids

Minimal media (MM) allows for the manipulation of individual medium components, but also generally requires that cells undertake extensive *de novo* synthesis that results in slower doubling times. To test if Mg^2+^ could modulate cell length in a defined medium, we utilized a phosphate-buffered glucose MM. We began with a base medium that contained 13 amino acids and 50.0 μM Mg^2+^ (MM-13aa). In MM-13aa, the doubling time for the *B. subtilis* 168 prototroph (functional *trpC+*) was 53 min with and without the addition of 10.0 mM MgCl^2^ (Fig 3A). Cell length in MM-13aa was assessed using microscopy and quantitation. As shown in Fig 3B and 3C, cells grown with supplemental Mg^2+^ were 45% shorter than those grown in the MM-13aa only.

**Figure 3.**
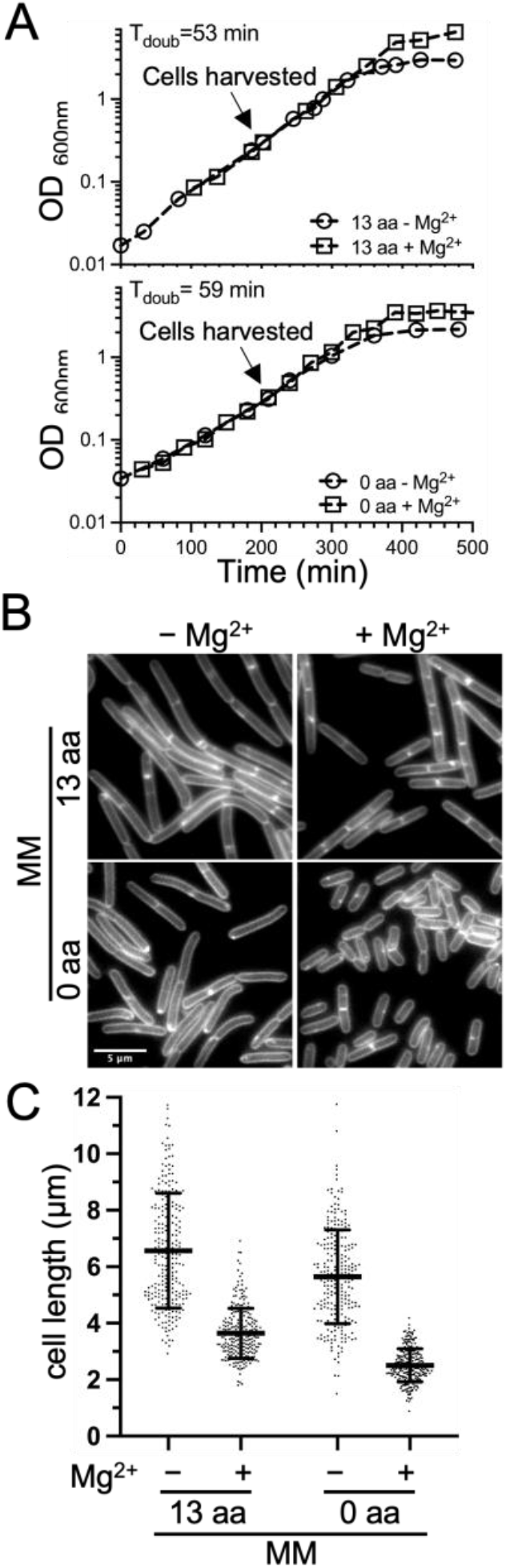
Cell length of *B. subtilis* following growth in minimal medium (MM). WT *B. subtilis* 168 (*trpC+*)(BTG169) was cultured in base MM containing 50.0 µM Mg^2+^ supplemented with 10.0 mM MgCl_2_ and amino acids when indicated (+ Mg^2+^). (A) Representative growth curves. (B) Micrographs following membrane staining and epifluorescent microscopy (scaled identically). (C) Scatter plots showing the distribution of cell lengths quantitated for 250 cells from each condition. Bars represent the means of 250 cells ± SD.

To test if the Mg^2+^-responsive phenotype was dependent on the presence of amino acids in the medium, the experiments were repeated in the base MM without amino acid supplementation. This modification increased the doubling time to 59 min, but the growth rate again remained unchanged with and without Mg^2+^ supplementation (Fig 3A). Similar to the pattern observed in the media that contained amino acids (LB, CH*, and MM-13aa), cells were shorter in the Mg^2+^ supplemented medium compared to the unsupplemented control (Fig 3B and 3C). These results suggest the Mg^2+^-responsive phenotype is generalizable to both complex and defined media and does not depend on the addition of amino acids.

### Mn^2+^ and Zn^2+^ supplementation also elicit cell shortening

We wondered if the growth-rate independent cell shortening was specific to Mg^2+^ or could also be induced by providing other metals in excess. For these experiments, we utilized the CH* medium, which contains 0.2 mM CaCl^2^, but otherwise only trace metals. Cells were grown as before, but instead of supplementing cultures with MgCl_2_, salts of Ca^2+^, Cu^2+^, Fe^2+^, Fe^3+^, Mn^2+^, and Zn^2+^ we added. Of the metals tested, only Mn^2+^ and Zn^2+^ produced significant cell shortening (Fig 4 and Fig S1); however, Zn^2+^ strongly reduced growth rate at the concentrations required to observe the cell shortening effect (0.2 mM)(Fig S1). While Mn^2+^ did not reduce the doubling time of cells once the culture reached exponential phase, the precultures containing Mn^2+^ remained in lag several hours longer than unsupplemented cells before OD_600_ increases were observed. Due to pleiotropic effects of Zn^2+^ and Mn^2+^ on growth, we chose to focus on Mg^2+^ for the remainder of the study.

**Figure 4.**
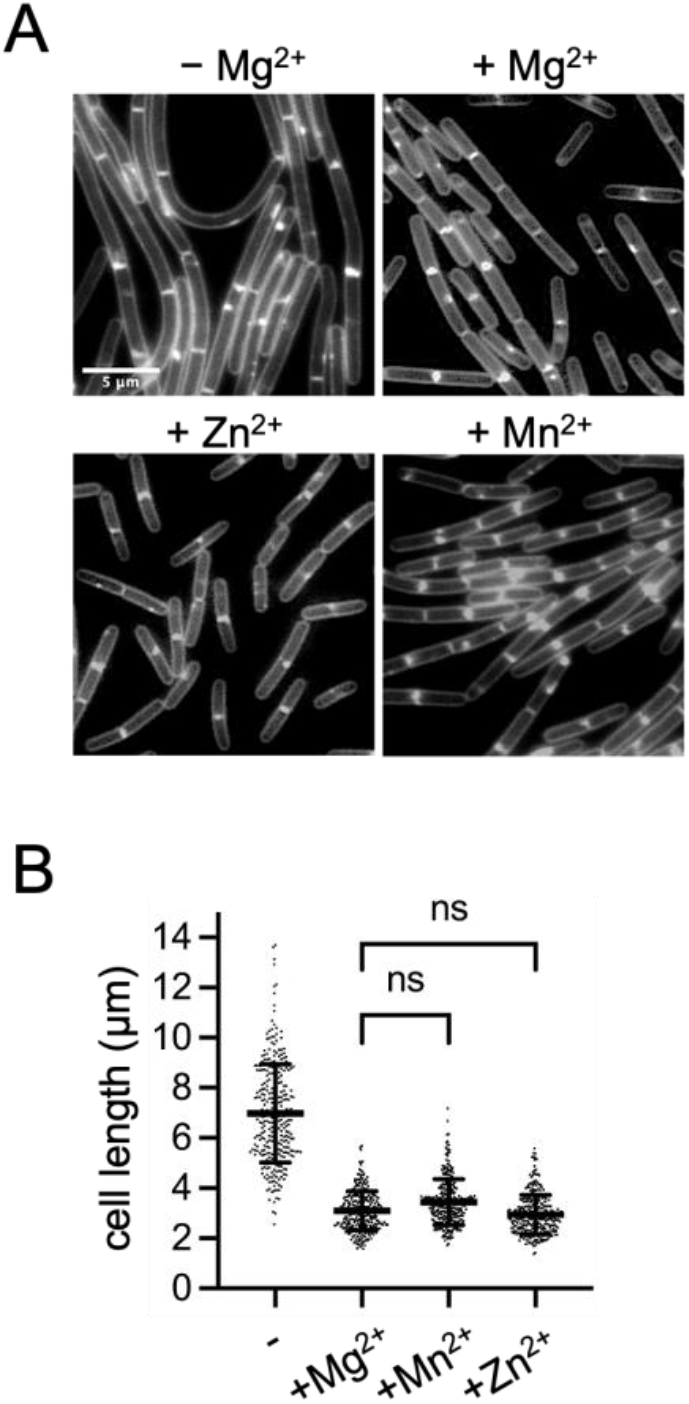
Effect of divalent metals on cell length. Wild-type 168 (BJH004) cells were cultured in CH* without or with the following concentrations of divalent cation salts: 0.1 mM MnSO_4;_ 0.2 mM ZnCl_2;_ 10.0 mM MgCl_2_. (A) Micrographs following membrane staining and epifluorescent microscopy (scaled identically). (B) Scatter plots showing the distribution of cell lengths quantitated for 300 cells from each condition. Bars represent the means of 300 cells ± SD. (ns, p>0.05).

### Mg^2+^-responsive cell length changes follow a dose-response curve

Next we wanted to know if cell length decreased proportionally with increasing extracellular Mg^2+^ or if, alternatively, there was a threshold at which cells underwent a switch in cell length. To test, we cultured cells in media across a range of Mg^2+^ concentrations, from 6.25 μM MgCl_2_ (where differences in growth rate begin to emerge, Fig 5A) up to 25.0 mM. Cells from each condition were imaged and the cell lengths were quantified (Fig 5). We found that cell shortening was neither directly proportional to Mg^2+^ availability, nor a switch. Instead, cells became progressively shorter from 12.5 μM to 4.0 mM, with the largest decrease occurring between 0.2 μM and 2.0 mM (Fig 5C and 5D).

**Figure 5.**
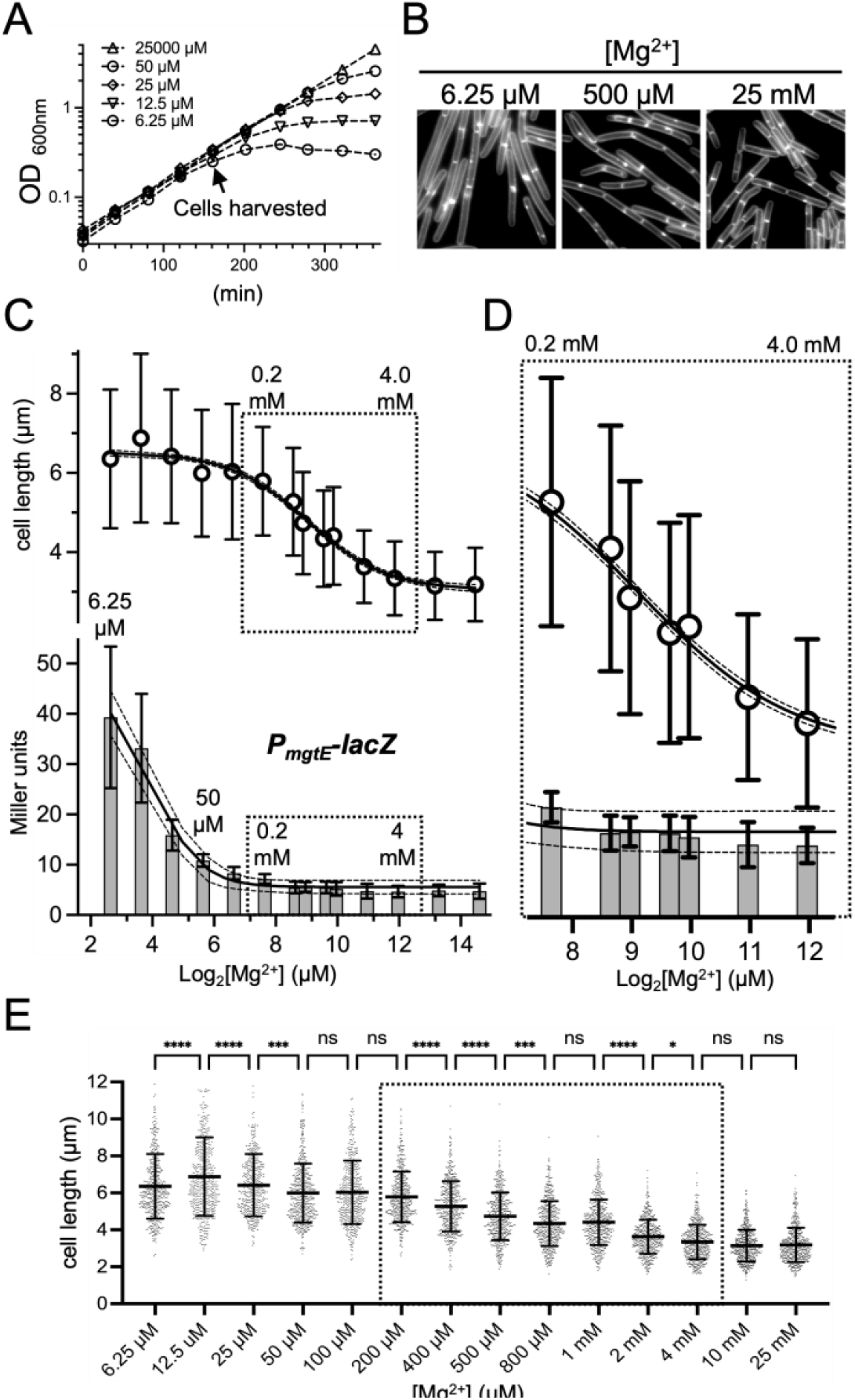
Correlation analysis of extracellular Mg^2+^ with cell length and a reporter of intracellular Mg^2+.^ WT cells harboring a reporter for *P*_*mgtE*_*-lacZ* (BTG182) were grown in MM-13aa with the indicated concentrations of supplemental MgCl_2_ (A) Representative growth curves and (B) representative micrographs following membrane staining (scaled identically) from across the range of Mg^2+^ concentrations examined. (C) Mean cell lengths for 500 cells ± SD for each condition (top) and β-galactosidase activity assay for the *mgtE* riboswitch transcriptional reporter (bottom). (D) Enlargement of regions boxed in graphs at left. The values above the SD bar indicate the concentration of MgCl_2_ added. (E) Scatter plots showing the distribution of cell lengths at each Mg^2+^ concentration. Bars represent the mean cell lengths for 500 cells ± SD. (****, p<=0.0001), (***, 0.0001<p<=0.001), (**, 0.001<p<=0.01), (*, 0.01<p<=0.05), (ns, p>0.05).

Cells in the 0.2 μM and 2.0 mM window grew at equivalent doubling times (Fig 5A), suggesting cells were not yet experiencing significant intracellular Mg^2+^ limitation. To independently assess, we introduced a promotor fusion (P_*mgtE*_-*lacZ*) that reports on changes in internal Mg^2+^ availability. The promoter region includes a riboswitch that terminates transcription when intracellular Mg^2+^ levels are sufficient (48, 49). Dann et al observed an approximately 100-fold induction of LacZ activity following extended growth of cells at 5.0 μM Mg^2+^ (growth-rate limiting) compared to Mg^2+^ excess (2.5 mM)(see Fig 1B in reference (48)). Consistent with these findings, we observed LacZ activity was modestly induced (∼10-fold) in media containing 6.25 μM Mg^2+^ (Fig 5B). Notably, the timepoint taken is just when growth rate effects begin to be observed for 6.25 μM Mg^2+^ (Fig 5A).

The steepest decline in cell-shortening we observed occurred between 0.4 and 1.0 mM. To our surprise, no difference in P_*mgtE*_-*lacZ* activity was detectable in this window, suggesting the cell shortening is unlikely to be attributable to increased levels of labile intracellular Mg^2+^. We considered the possibility that the riboswitch of the P_*mgtE*_-*lacZ* reporter may already be saturated at 0.4 mM external Mg^2+^ and thus insensitive to further increases. To test, we deleted the gene for the Mg^2+^ exporter protein MpfA, previously shown to result in an increase in intracellular Mg^2+^ (50). Even when cells were grown in 10.0 mM MgCl_2_, the Δ*mpfA* mutant displayed an additional 4-fold reduction in LacZ activity compared to wildtype (Fig 6A), indicating that the reporter retained sensitivity. This result suggests that if there are changes in intracellular Mg^2+^ in the 0.2 to 1.0 mM range where we observe the most dramatic decreases in cell length, then the changes are too small to be detected with our reporter. Notably, even though intracellular Mg^2+^ apparently increased in the Δ*mpfA* mutant compared to wildtype, no differences in average cell length were observed (Fig 6B and 6C). Collectively, these results suggest that under our growth conditions, changes in intracellular labile Mg^2+^ are neither necessary, nor sufficient to elicit cell shortening. At the same time, we do not exclude the possibility that the cells may be responding to changes in intracellular Mg^2+^ not detectable by P_*mgtE*_-*lacZ*, such as an increase in the non-labile pool. An alternative possibility is that the response is driven by extracellular, or at least extracytoplasmic, Mg^2+^.

**Figure 6.**
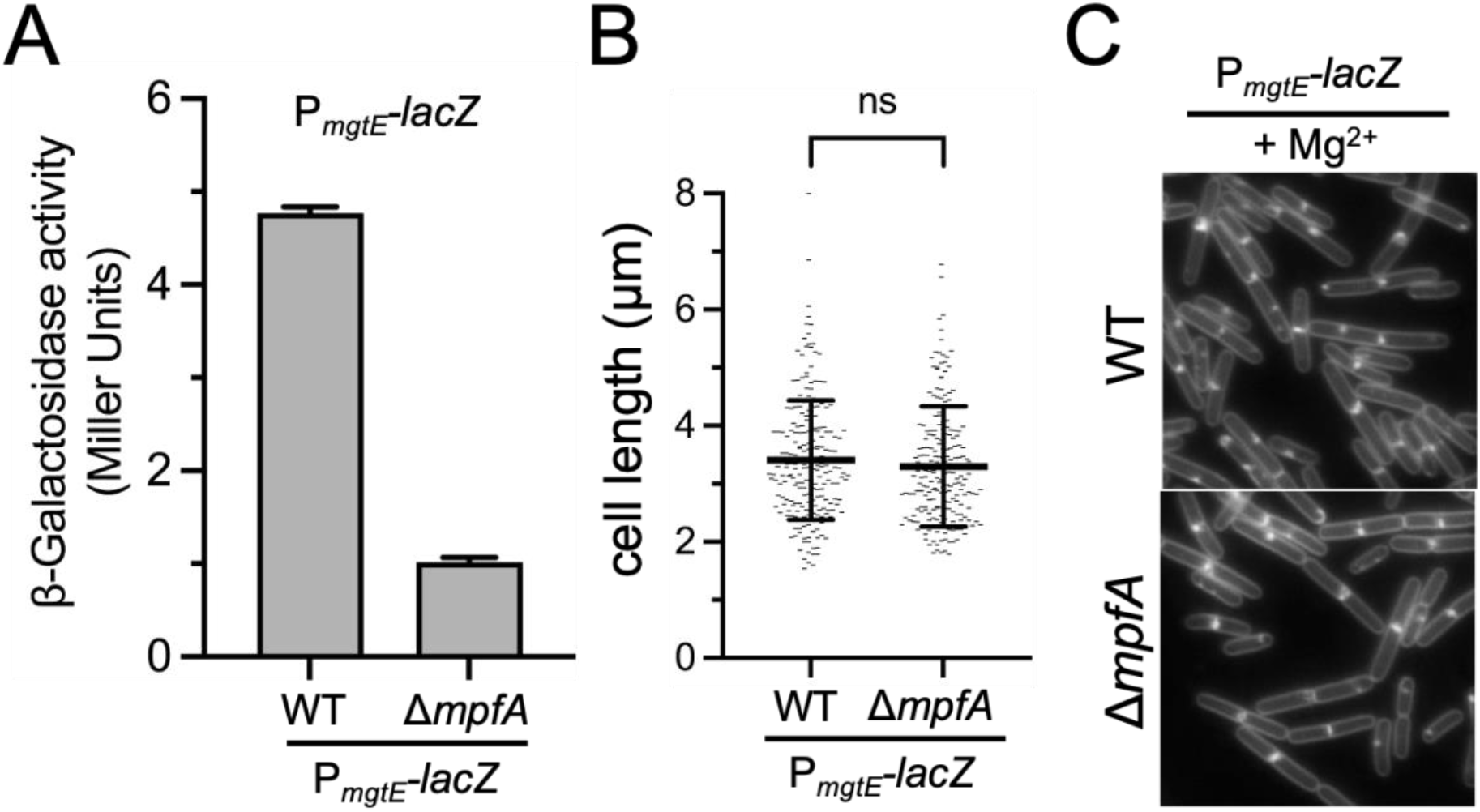
Monitoring expression of an intracellular reporter of Mg^2+^ and cell length in a Mg^2+^ exporter mutant background (Δ*mpfA*). WT (BTG182) or the Δ*mpfA* mutant (BTG333) harboring a reporter for P_*mgtE*_*-lacZ* were grown in MM (13 aa) supplemented with 10.0 mM MgCl_2_. Samples were taken during exponential growth. (A) β-galactosidase activity assay. Bars represent means from 2 biological replicates ± SD. (B) Scatter plots showing the distribution of cell lengths. Bars represent the mean cell lengths for 200 cells ± SD. (ns, p>0.05). (C) Representative micrographs of cells stained with TMA and scaled identically.

### Cell shortening following a shift from lower to higher Mg^2+^ is rapid

In the experiments above, the cell lengths were determined during steady-state growth, taking care to collect cells at equivalent densities because Mg^2+^ continues to be depleted from the media with time. To assess the transition from longer to shorter, we grew cells in two different base media and monitored cell length at 10 min intervals after the addition of 10.0 mM MgCl_2_ (Fig 7). For the media tested, the mean cell length continued to decrease for 40-50 min, at which point cells achieved a mean length that was similar to cells always grown with excess Mg^2+^ (Fig 7). Remarkably, 54% and 40% of the total decrease in mean cell length occurred within the first 10 min of adding Mg^2+^ in the CH* and MM-13aa mediums, respectively (Fig 7). Thus, the initial response to Mg^2+^ is relatively rapid, well below the doubling time of the cells. After the initial rapid decrease, a more gradual decline in average cell length was observed. These results suggest the adjustment to higher Mg^2+^ may occur through two distinct mechanisms; one that is immediately triggered following the addition of Mg^2+^, and one that requires an outgrowth period of more than a generation time before a new steady-state length is achieved.

**Figure 7.**
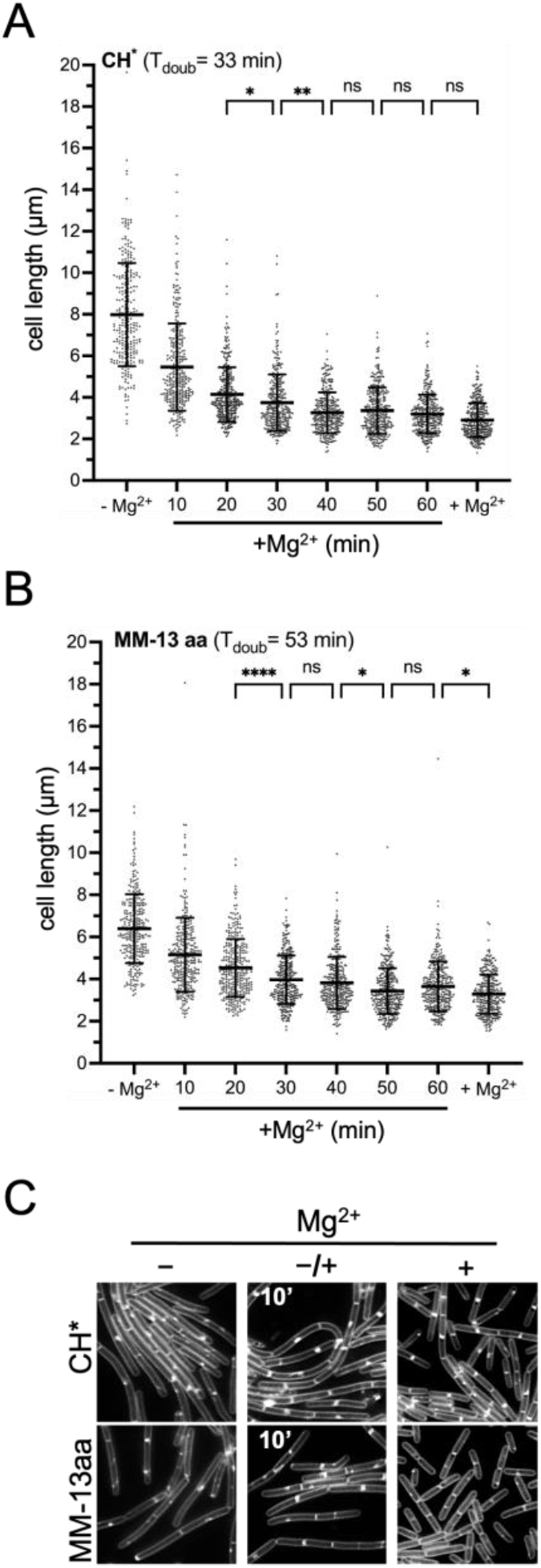
Monitoring *B. subtilis* cell length changes following Mg^2+^ supplementation. Wild-type 168 (BJH004) was cultured in the indicated media without Mg^2+^ supplementation (-Mg^2+^). Following addition of 10.0 mM MgCl_2_, cells were imaged at 10 min intervals. (A and B) Bars represent the mean cell lengths for 300 cells ± SD. (****, p<=0.0001), (**, 0.001<p<=0.01), (*, 0.01<p<=0.05), (ns, p>0.05). (C) Representative micrographs. Membranes are stained with TMA and images are scaled identically.

### Mg^2+^ modulates the frequency of Z-ring assembly

A rod-shaped bacterium can increase cell length without altering other dimensions by elongating faster, dividing less frequently, or both. Based on optical density, the growth rate of the cells in our experiments was equivalent with and without Mg^2+^ supplementation. Although absorbance readings are the most widely accepted method for monitoring cell growth, we considered the possibility that the populations we were comparing might absorb light differently enough to mask differences in mass accumulation. To independently test, we examined the abundance of a constitutively expressed protein, SigA, using western blot analysis. In both CH* and MM-13aa, SigA levels were equivalent with and without Mg^2+^ supplementation when cells were normalized to each other using OD_600_ values (Fig S2A). As an additional control, we compared the dry weight of samples grown in MM-13aa with and without excess Mg^2+^ (Fig S2B). The dry weights were indistinguishable in the two conditions, further supporting the conclusion that optical density provides an accurate approximation of cell mass. These results strongly support the conclusion that Mg^2+^ supplemented cells become shorter as a result of more frequent cell divisions, and conversely, that cells divide less often in the lower Mg^2+^ conditions.

A delay in septation could occur before or after assembly of the divisome. To assess, we grew wildtype harboring a GFP fusion to ZapA, an early-arriving cell division protein that colocalizes with FtsZ as part of the so-called “Z-ring” (51, 52). Cells expressing P_*xylA*_*-GFP*-*zapA* grew equivalently with and without 10.0 mM MgCl_2_ (Fig 8A) and retained the Mg^2+^-responsive reduction in cell length (Fig 8B and 8C). Epifluorescence microscopy was performed, and images were captured of membranes, DNA (nucleoids), and Z-rings from both conditions (Fig 8B). Overlays of the micrographs were used to quantitate both the number of distinct nucleoids per cell (Fig 8D) and the number of Z-rings per unit cell length (Fig 8E).

**Figure 8.**
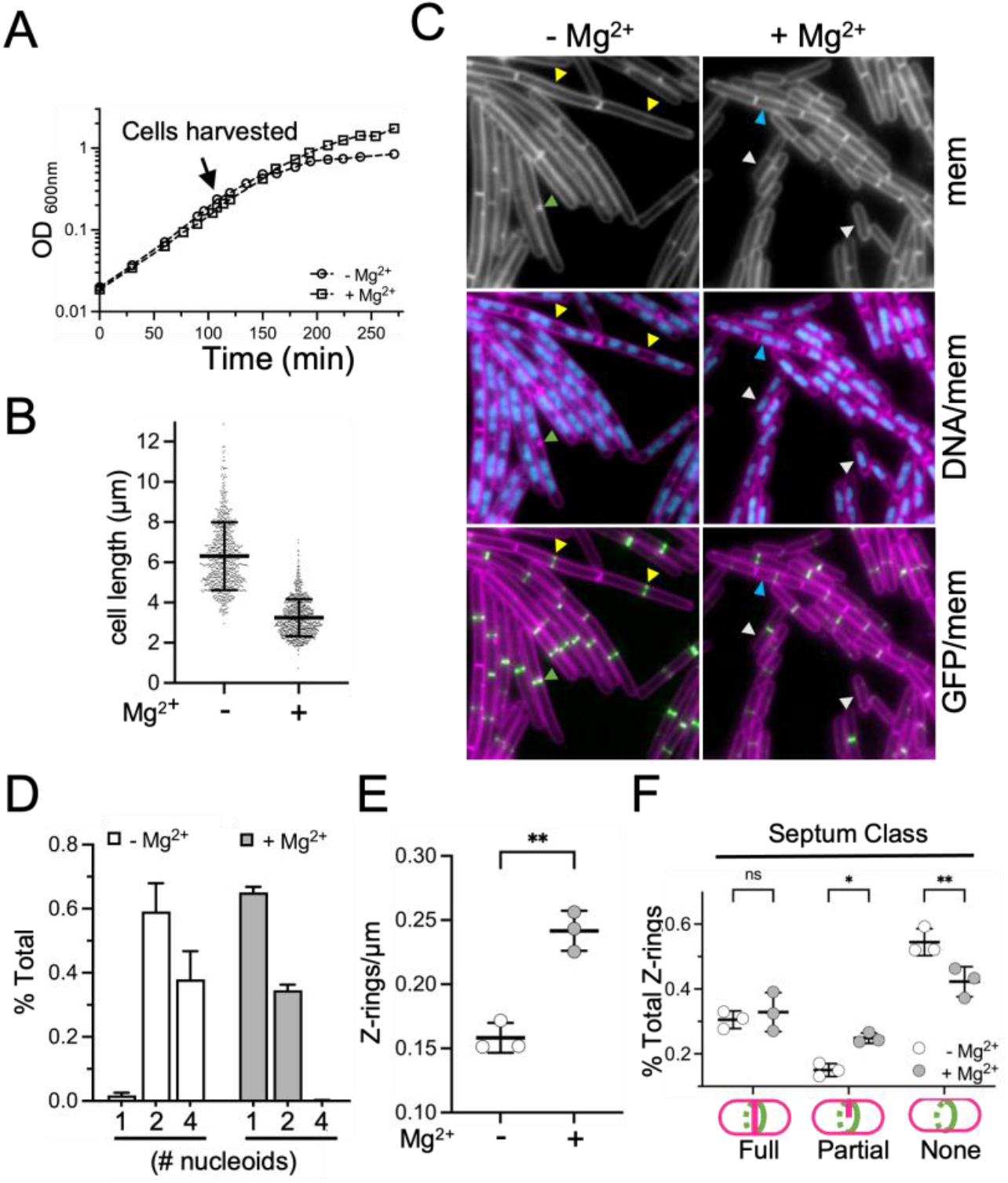
Mg^2+^ modulates the frequency of Z-ring assembly. WT cells harboring *P*_*xyl*_*-GFP*-*zapA* (BTG186) were cultured in CH* with 2.0 mM xylose. 10.0 mM MgCl_2_ was added as indicated. (A) Representative growth curves. (B) Scatter plots showing the distribution of cell lengths. Bars represent the mean cell lengths 800 cells ± SD. (C) Representative micrographs following staining of membranes and nucleoids with FM4-64 and DAPI, respectively. Images are scaled identically. Arrowheads indicate examples of cells with four nucleoid masses and Z-rings, that lack septa (yellow), one nucleoid mass (white), two nucleoid masses (blue), or partial septa (green). (D) Fraction of cells with the indicated number of nucleoids from three independent biological replicates. (E) Average number of Z-rings per unit cell length ± SD. Each circle represents the mean of 300 cells from three independent biological replicates per condition. (**, 0.001<p<=0.01). (F) Average fraction of cells with coalesced ZapA-GFP presenting the indicated septum type ± SD. Each circle represents the mean of 300 cells from three independent biological replicates per condition. (**, 0.001<p<=0.01), (*, 0.01<p<=0.05), (ns, p>0.05).

Regardless of condition, both populations possessed a large proportion of cells with two nucleoids (35% and 59% for with and without supplemental Mg^2+^, respectively). The most striking differences became apparent when comparing the proportion of cells with one or four nucleoids. While only 2% of cells grown in CH* had one distinct nucleoid mass, this proportion increased 30-fold (to 65%) in the Mg^2+^-supplemented medium. Conversely, cells with four nucleoids were relatively frequent (38%) in the base medium, but rare (<1%) in the Mg^2+^-supplemented cultures (Fig 8D). These results indicate division is more frequent when Mg^2+^ is in excess. Consistent with this idea, Z-rings were more frequently observed along the length of Mg^2+^-supplemented cells and fewer Z-rings were observed along the length of cells grown in unsupplemented CH* (Fig 8E); moreover, Z-rings that did form were less likely to be associated with a partial (in-process) septation (Fig 8F). Similar results are observed when FtsZ was tracked directly with an FtsZ-GFP fusion (Fig S3). We conclude that there is a window of Mg^2+^ availability that can modulate cell division frequency before growth rate is impacted.

### Overexpressing undecaprenyl pyrophosphate synthetase (uppS) results in loss of the Mg^2+^-responsive phenotype and constitutively short cells

In the CH* shifting experiment, more than half of the decrease in average cell length occurred within 10 min of adding Mg^2+^ (Tdoub = 33 min)(Fig 7), suggesting a majority of cells were generally poised to divide, but somehow inhibited. In a 1969 study, Garrett showed that Mg^2+^ limitation caused *B. subtilis* (W23) suspensions to accumulate the cytoplasmic PG precursor UDP-MurNAc-pentapeptide (53). This result suggests that the next step in the PG synthesis pathway, generation of Lipid I by conjugation of MurNAc-pentapeptide to Und-P (Fig S4A), is sensitive to Mg^2+^ availability.

Und-P is generated through dephosphorylation of Und-PP, which is either regenerated following transfer of Lipid II proteoglycan subunits to the PG layer or synthesized de novo by UppS (Fig S4A). To test if increasing Und-PP pools from the de novo pathway would promote more frequent division, we overexpressed *uppS* from an IPTG-inducible promoter. We found *uppS* overexpression resulted in cells that were equivalently short, irrespective of Mg^2+^ supplementation (Fig 9A and 9B). By contrast, cells overexpressing *mraY*, which encodes the enzyme that reversibly transfers phospho-MurNAc-pentapeptide to Und-P (FigS4A)(54), were still responsive to Mg^2+^ (Fig S4B), while those overexpressing *bcrC* (encoding the phosphatase that generates Und-P from Und-PP (Fig S4A)) were not (Fig S4C and D). Together these results suggest that availability of Lipid I, but not phospho-MurNAc-pentapeptide, is rate-limiting for cell division in the unsupplemented media. The growth rate of cells overexpressing *uppS* was equivalent with and without Mg^2+^ supplementation (Fig 9C), so the smaller cells is not simply attributable to the slower growth rate. At the same time, we cannot exclude the possibility that the slow growth reduces cells to a length that makes differences too subtle to be measured with our assay.

**Figure 9.**
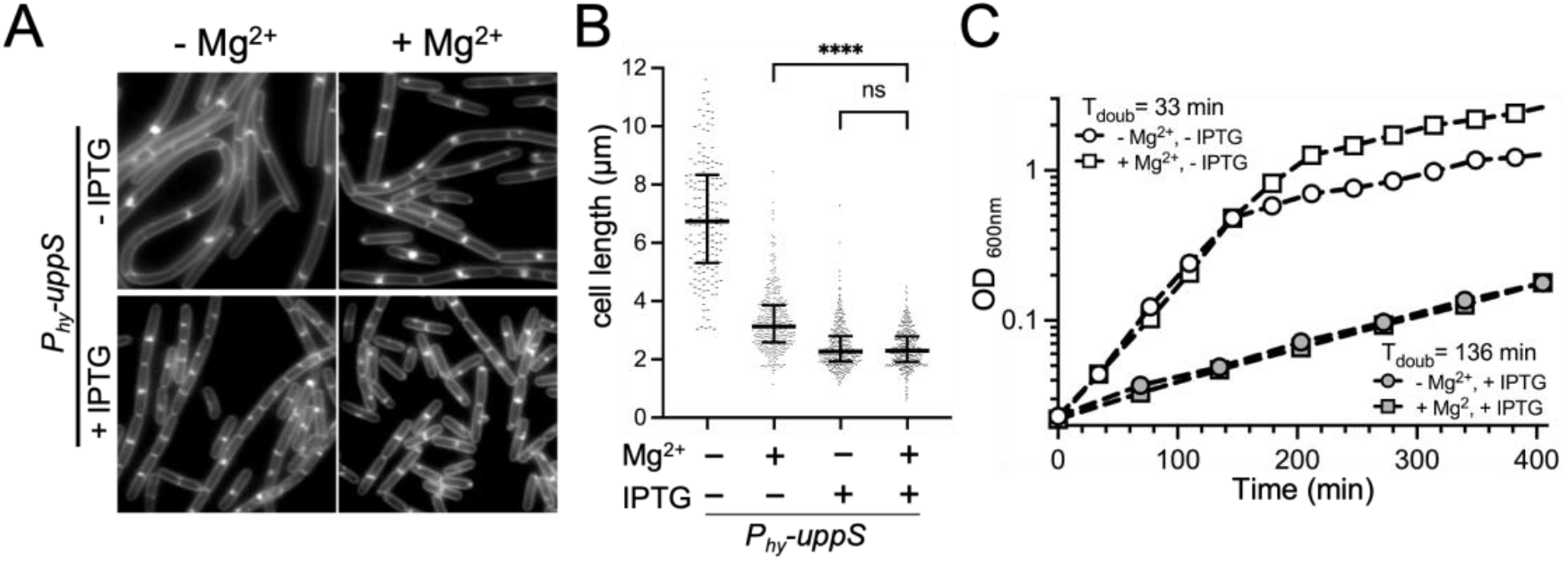
UppS overexpression results in constitutively short cells. WT cells harboring *P*_*hy*_*-uppS* (BTG708) were cultured in CH* with MgCl_2_ (10.0 mM) and IPTG (0.5 mM) added as indicated. (A) Representative micrographs of cells stained with TMA, scaled identically. (B) Scatter plots showing the distribution of cell lengths quantitated for 300 cells from each condition. Bars represent the mean cell length ± SD (Ordinary one-way ANOVA). (****)p<0.0001, (ns)p>0.01234. (C) Representative growth curves.

### RNA-seq analysis of cells grown with and without excess Mg^2+^

To explore the Mg^2+^-responsive phenotype further, we used RNA-seq analysis to determine if there were relative changes in transcription between the short and long cells that could provide insight into mechanism. CH*-grown cells were treated as the control and CH* with 10.0 mM MgCl_2_ as the experimental. Genes expressed more or less in the Mg^2+^-supplemented medium relative to CH* only were categorized based on known regulatory information retrieved from SubtiWiki (Table S1 and S2)(55).

As expected based on the riboswitch data (Fig 5), transcriptional changes related to Mg^2+^ homeostasis were not significant. To our surprise, a number of other regulons related to metal homeostasis showed significant and internally consistent relative shifts in transcription. In particular, the profiles suggest upregulation of genes repressed by Fur (Fe^2+^ acquisition, sequestration, and sparing), MntR (Mn^2+^ uptake), and Zur (Zn^2+^ acquisition). In addition, there was enhanced expression of a large number of genes consistent with a response to enhanced general and oxidative stress (SigB, PerR). A subset of genes from two different prophage (PBSX and SPβ) were also expressed at higher levels, but not in a manner consistent with prophage induction (Table S1).

Conversely, and consistent with the observed up-regulated genes, we observed downregulation of the CzrA repressed genes *cadA, czcD*, and *trkA (*Zn^2+^ efflux/resistance to toxic metals), *mneP* (MntR-activated Mn^2+^ efflux), a large number of genes under control of various cell envelope stress sigma factors (SigM/X/W/V), and those activated by the two-component response regulator YvrHb (56). Notably, several regulatory genes showed decreased transcription including *rsiX* (encoding a SigX anti-sigma factor), *sigX*, and *abh* (a regulator of transition state).

## DISCUSSION

The findings in this study surprised us in a number of ways. First, the discovery of the cell division phenotype was itself unanticipated. Although LB is known to be low in Mg^2+^ and is generally discouraged by cell physiologists for various reasons (nicely outlined in a Small Things Considered blogpost by Hiroshi Nikaido (57)), LB has also been the default medium for routine propagation of *B. subtilis* and *E. coli* for decades; not necessarily a context in which one expects to observe a novel phenotype. We cannot definitively say whether or not the phenotype observed in LB (Fig 1) was due to unusually low Mg^2+^ in the batch, as the powder was exhausted, and its bottle along with its lot number discarded long before we appreciated the phenotype’s significance. Nonetheless, the data strongly implicated Mg^2+^ and motivated further experiments ultimately demonstrating that reduced Mg^2+^ availability impacts division before growth rate and cell elongation.

The finding itself makes sense intuitively, as cell division is not a prerequisite for reproduction of genetic material, growth, or persistence. In fact, many bacteria are known to subsist as filamentous forms that divide infrequently. An extreme version of the filamentous lifestyle was recently discovered in the marine organism *Thiomargarita magnifica*, which average lengths of nearly a centimeter (58). This remarkable bacterium was found to harbor ∼37,000 copies of its genome per millimeter of cell length, evidence of a dramatic uncoupling between growth and cell division. It is notable that many of the longest bacteria ever recorded, including the giant bacterium *Beggiatoa* discovered by Winogradsky, are sulfur-oxidizers (59). Evidence for an intimate relationship between sulfur metabolism and cell division is not restricted to giant bacteria. *E. coli* mutants with reduced ability to convert ATP and the sulfur-containing amino acid methionine into S-adenosyl methionine (SAM) are inhibited for cell division but continue to grow and replicate DNA (60). The capacity of cells to uncouple division from other biosynthetic processes makes sense from an evolutionary perspective because it affords cells a mechanism to generate new “units” even as resources become depleted while simultaneously preserving the capacity to separate into individual cells under more favorable conditions.

Another surprising result was the degree and type of transcriptional changes that occurred between the supplemented and unsupplemented growth conditions. We expected that if changes occurred at all, then they would be associated with envelope synthesis or Mg^2+^ uptake or efflux. Instead, we observed changes in genes related to general stress response and Fe^2+^, Mn^2+^, and Zn^2+^ homeostasis. More specifically, we found that when Mg^2+^ is more available, expression of the SigB regulon is enhanced, and the Fur, MntR, and Zur regulons undergo shifts consistent with acquisition, sequestration, and/or sparing responses.

At first, the metal responses were perplexing as we are accustomed to thinking about these metal regulons in terms of responses to starvation or toxicity, conditions not present in our experiments. However, the responses are relative and not absolute, and we can think of several other ways to interpret the data. The first is to think of the changes as homeostatic adjustments. For example, the data indicate that in the higher Mg^2+^ condition the cells receive signals consistent with the need to acquire, sequester, and spare Fe^2+^, and increase internal Mn^2+^ and Zn^2+^. Expression of genes for surfactin biosynthesis (*srfABCD*), and the SigB general stress response are also increased. Fe^2+^ sequestration and Fe^2+^-sparing responses would reduce the possibility of Fenton reactions (61), Mn^2+^ and Zn^2+^ both reduce internal reactive oxygen species (62-68), and surfactin and SigB reduce oxidative/energy stress, the former by reducing proton motive force (69, 70). The overall profile suggests that as an averaged population, the Mg^2+^-supplemented cells experience signals indicating a response to oxidative stress than unsupplemented cells.

The second possibility, which is not mutually exclusive, is that at least some of the transcriptional changes are associated with events just before, after, and/or during cell division itself. Cell-cycle variations are typically obscured when samples are collected from pooled populations of asynchronously growing cells. The samples collected for RNA-seq were also growing asynchronously; however, because the Mg^2+^-supplemented populations were essentially “enriched” for dividing cells (Fig 8 and Fig S3), transcriptional changes associated with division should also be enriched. The idea that organisms may experience different stress signals and adjust metal pools not only to respond to stress or environment, but also as regulated part of the life cycle or development is intriguing, and we think merits further inquiry.

We noted one anomaly in the RNA-seq data that we do not understand. In the Zur regulon, we see more RNA corresponding to *yczL*, but relatively little or no increase in other genes found in the predicted (71) *folEB*-*yciB*-*yczL*-*zagA* transcript (Table S1). The unexpected differential expression of *yczL* region could be attributable to differences in RNA stability rather than transcription itself. The gene upstream of *yczL* (*yciB*), is proceeded by four instances of putative co-translational coupling, the last of which could lead to translation of *yczL*. It may be worth exploring if there is functional consequence to the predicted coupling, such as altering transcriptional readthrough, transcript stability, or expression of the downstream gene, *zagA*.

We do not know if any of the transcriptional changes relate directly to the Mg^2+^-responsive phenotype or are incidental. Would differential expression still occur if were grown at maximal “shortness” but different Mg^2+^ concentrations (4.0 mM-25.0 mM range, Fig 5)? The cell shortening phenotype was still detected in a prophage-cured strain (Table S3), so at least these changes appear to be incidental. Consistent with this conclusion, others have documented expression from various *B. subtilis* prophage loci under different growth regimes (72).

Visually screening a number of deletion strains also failed to implicate any single gene knockouts in the Mg^2+^-responsive phenotype (Table S1-S3). The only condition identified that showed an apparent loss of Mg^2+^-responsiveness in CH* was overexpression of *uppS* (Fig 9). However, as these cells grew slowly we could not confidently exclude the possibility that 1) cell shortening occurred but fell outside the detection limit of our assay and/or 2) the slow growth rate imposed an upper limit on division frequency that was dominate to the effects of Mg^2+^. Still, it is plausible that exogenous Mg^2+^ somehow enhances availability of Und-P to the divisome. In the shifting experiment (Fig 7), we observe two paths to cell shortening; a dramatic initial decrease in cell length resulting from rapid division onset, and a second more protracted period where average cell length decreases more incrementally. These observations suggest that the initial response may have a more biophysical or enzymatic basis, while the second may require biosynthesis and outgrowth to achieve a new steady-state. One idea that is purely speculative, is that the rapid division induced by the sudden switch to higher Mg^2+^ is driven by more rapid flipping of Und-P to the cytoplasmic face of the membrane and/or enhanced conversion of Und-PP to Und-P.

In summary, we show that the frequency of cell division can be uncoupled from growth rate by manipulating a single nutrient, Mg^2+^. Rich media are convenient, but have the weakness of variable formulation with regard to absolute and relative amounts of cofactors, trace metals and amino acids. Not only can this lead to confusing and irreproducible results, but also (as we unintentionally discovered), rich media may be masking some important biology.

## MATERIALS and METHODS

### General methods

Strains and details of strain construction can be found in the supplementary materials (Table S4 and Text S1). Cells were stored at -80°C in 15% glycerol (v/v). Strains were streaked for isolation on Lysogeny Broth, Lennox (LB) containing bactoagar (1.5% w/v) and incubated overnight (∼16 hr) at 37°C. Cultures for experiments were begun with single colonies from same-day plates by inoculating single colonies into a 20 mm glass tube containing 5 mL of media. The tube was incubated at 37°C in a roller drum until exponential stage. For experiments, 25 ml of medium in a 250 ml baffled flask was inoculated with exponential stage cultures (see above) and cells were incubated in a shaking water bath set to 280 rpm and 37°C.

LB was made by dissolving 20 g of Difco™ LB Broth, Lennox (product #240230) in 1 L ddH_2_O, followed by sterilization in an autoclave. CH* (1L) contained 10.0 g Casein acid hydrolysate (Acumedia, Lot No.104,442B), 3.7 g L-glutamic acid (25.0 mM), 1.6 g L-asparagine monohydrate (10.0 mM), 1.25 g L-alanine (14.0 mM), 1.36 g (10.0 mM) KH_2_PO_4_ anhydrous, 1.34 g (25.0 mM) NH_4_Cl, 0.11 g Na_2_SO_4_ (0.77 mM), 0.1 g NH_4_NO_3_(1.25 mM), 0.001 g FeCl_3_•6H_2_O (3.7 mM) and ddH_2_O. The pH was adjusted to 7.0 with 10.0 N NaOH before media was sterilized in an autoclave. After autoclaving, CaCl_2_ was added to 0.2 mM and L-Tryptophan was added to 0.1 mM. The base minimal medium (MM) contained 15.0 mM (NH_4_)_2_SO_4_, 5.0 mM KH_2_PO_4_, 50.0 mM Tris-HCl [pH 7.5], 27.0 mM KCl, 0.05 mM FeCl_3_, 0.01 mM MnSO_2_, 0.01 mM ZnSO_4_, 0.2% glucose (w/v), 27.0 mM sodium citrate, and 10.2 mM CaCl_2_. 50.0 µM MgCl_2_ was added to the to create the base medium used in MM experiments except for those in Fig 5, which required final concentrations below 50.0 µM. In these experiments, MgCl_2_ was added to the concentrations indicated in the figure. For MM-13aa, 10X filter-sterilized amino acids [pH 7.5] were added to 1X final concentration. The 10X mixture consisted of the following concentrations of L-amino acids: 250.0 mM Glutamate; 100.0 mM each of Alanine, Asparagine, and Proline; 50.0 mM each of Aspartic acid, Phenylalanine, Glycine, Isoleucine, Leucine, Threonine, and Valine; 10.0 mM each of Histidine and Tryptophan. The amino acids Arginine, Cysteine, Glutamine, Lysine, Methionine, Serine, and Tyrosine were excluded.

### β-galactosidase assays

*B. subtilis* strains were grown 250 mL baffled flasks with 25 ml of the indicated medium at 37°C and 280 rpm to the target OD_600_. The optical density was recorded and 1 mL of culture was harvested by 21,130 X g for 1 min at room temperature, removing supernatant by aspiration. Pellets were immediately stored at -80°C. To perform the assay, pellets were thawed, re-suspended in 0.5 mL Z-buffer (40.0 mM NaH_2_PO_4_, 60.0 mM Na_2_HPO_4_, 1.0 mM MgSO_4_, 10.0 mM KCl, 40.0 mM β-mercaptoethanol and 0.2 mg/ml lysozyme) and incubated at 30°C for 15 min. 100 µL of 4.0 mg/mL 2-nitrophenyl-β-D-galactopyranoside (ONPG) in Z-buffer was added and the samples were incubated at 30°C until pale yellow. The reaction was stopped with 250 µL 1.0 M Na_2_CO_3_ and the reaction time was recorded. The sample was vortexed for 5 sec, centrifuged at 21,130 X g for 3 min at room temperature in a tabletop centrifuge. The supernatant (minimum volume 0.8 mL) was transferred to a 1 ml cuvette and the OD_420_ and OD_550_ absorbance readings were recorded. β-galactosidase specific activity in Miller Units was calculated using the following formula: [OD_420_ -(1.75 x OD_550_)] / (time [min] x volume x OD_600_) x 1000.

### Western blot analysis

One mL of culture was collected and spun at 21,130 X g for 1 min at room temperature at the exponential stage and the OD_600_ value at time of sampling was recorded. the pellet was re-suspended in the lysis buffer (20.0 mM Tris [pH 7.5], 10.0 mM EDTA, 1 mg/mL lysozyme, 10 µg/mL DNase I, 100 µg/mL RNase A, 1.0 mM PMSF, 1 µL protease inhibitor cocktail (Sigma P8465-5ML) resuspended in 1 mL lysis buffer) to give a final OD_600_ equivalent of 15. The samples were incubated at 37°C for 10 min followed by addition of an equal volume of sodium dodecyl sulfate (SDS) sample buffer (0.25 M Tris [pH 6.8], 4% (w/v) SDS, 20% (w/v), 20% glycerol (v/v), 10.0 mM EDTA, and 10% β-mercaptoethanol (v/v)). Samples were heated for 5 min at 100°C prior to loading. Proteins were separated on 12% SDS-PAGE polyacrylamide gels, transferred to a nitrocellulose membrane at 100V for 60 min, and then blocked in 1X PBS containing 0.05% (v/v) Tween-20 and 5% (w/v) dry milk powder. The blocked membranes were probed overnight at 4°C with anti-SigA (1:20,000, rabbit, gift from Fujita Masaya, University of Houston, Houston, TX) diluted in 1X PBS with 0.05% (v/v) Tween-20 and 5% (w/v) milk powder). The membranes were washed three times with 1X PBS containing 0.05% (v/v) Tween-20 and transferred to 1X PBS with 0.05% (v/v) Tween-20 and 5% (w/v) milk powder containing 1:5,000 horseradish peroxidase-conjugated goat anti-mouse IgG secondary antibody (AbCam ab205719) and incubated on a shaking platform for 1 hr at room temperature. The membranes were washed 3X with 1X PBS containing 0.05% (v/v) Tween-20 and signal was detected using SuperSignal West Femto maximum sensitivity substrate (Thermofisher) and a Biorad Gel Doc Imaging System.

### Fluorescence microscopy

1mL cultured cells at exponential stage were harvested and concentrated by centrifugation at 6,010 X g for 1 min and re-suspended in 5 µL 1 X PBS with either 1-(4-(trimethylamino)phenyl)-6-phenylhexa-1,3,5-triene (TMA-DPH)(50.0 µM) or N-(3-Triethylammoniumpropyl)-4-(6-(4-(Diethylamino) Phenyl) Hexatrienyl) Pyridinium Dibromide (FM4-64)(6 µg/mL) and 4′,6-diamidino-2-phenylindole (DAPI)(2 µg/mL). All dyes were purchased from ThermoFisher. For the Zap-GFP and FtsZ-GFP experiments, cells were mounted on 1% (w/v) agarose pads made with PBS [pH 7.4] and overlaid with an untreated glass coverslip. Otherwise, cells were mounted on glass sides with polylysine-treated coverslips prior to imaging.

Cells were imaged on a Nikon Ti-E microscope using a CFI Plan Apo lambda DM 100X objective and X-Cite XYLIS 365nm illumination system (Excelitas Technologies). Filter cubes utilized were C-FL UV-2E/C (DAPI), C-FL Texas Red HC HISN Zero Shift, and GFP HC HISN Zero Shift. Micrographs were acquired with a CoolSNAP HQ2 monochrome camera using NIS Elements Advanced Research and analyzed in ImageJ (73).

### Cell length measurements

TMA-DPH micrographs were analyzed with ImageJ (73). For each cell, a line was placed the extended from pole-to-pole. One compartment between two bright solid septa with homogenous signals were counted as one intact cell. Cells with bright partial septa or septa interrupted with discrete dark areas were counted as cells with partial septa.

### Quantitation of Z-rings per unit cell length

The cell length line (see above) was overlapped with the GFP micrograph of the Z-ring and the number of Z-rings along the cell length of the line was counted. When two cells shared a pole and a Z-ring, each Z-ring was counted as 0.5/cell. The Z-ring number per µm was calculated by dividing the Z-ring total number by the total cell length.

### Nucleoid number per cell

The cell length line was overlapped with the DAPI micrograph and cells were classified based on the number of nucleoids overlapping the length label. The number of nucleoids per cell was calculated by dividing the number of cells containing each number of nucleoids (1, 2, or 4) by the total number of cells counted.

### Classification of septa in cells containing coalesced Z-rings

Micrographs of the GFP channel (ZapA-GFP or FtsZ-GFP) were overlaid with the corresponding TMA-DPH micrograph. The cells were classified as having no, partial, or full septa. The fraction of each septum type was determined by dividing the number of cells in each class by the total number of cells counted.

### Statistical analysis and data plotting

Graphs were generated and statistical analysis was performed using GraphPad Prism version 9.4.0 for Mac (GraphPad Software, San Diego, California USA, www.graphpad.com). Statistical analysis in the following figures were performed with One-way ANOVA followed by Tukey’s multiple comparisons test, assuming Gaussian distribution with equal SDs. (Fig 4B, Fig 5E, Fig 7A&B, Fig S2B, Fig S3C, Fig 9B) Statistical analysis in the following figures were performed with Two-way ANOVA followed by Tukey’s multiple comparisons test, fitting a full model. (Fig 8F, Fig S3D) Statistical analysis in the following figures were performed with Two-tailed unpaired t-test, assuming Gaussian distribution and both populations have the same SD. (Fig 1B, Fig 2D, Fig 6B, Fig 8E).

### RNA-seq

Samples were collected from three independent biological samples. Cells were cultured in CH* medium with and without 10.0 mM MgCl_2_ and 0.5 mL cells were collected at exponential stage (OD_600_ of approximately 0.2) by centrifuging at 21,130 X g for 2 mins at room temperature. RNA was collected using RNAprotect Bacteria reagent and RNeasy mini kit (Qiagen) according to the manufacturer’s instructions. RNA sequencing was performed by the Texas A&M AgriLife Research Genomics and Bioinformatics Service (College Station). Total RNA were prepared for sequencing using TruSeq(tm) Stranded Total RNA and Ribo-Zero Gold (Illumina). RNA-seq was performed using the HISeq 4000 platform (Illumina). RNA-seq data was processed through HISAT2 (74) and StringTie (75). Differential expression analysis was done using the Deseq2 R package (76). Regulons were assigned using the GinTool starter package obtained from Dr. L.W. Hamoen (University of Amsterdam, Swammerdam Institute for Life Sciences). The cutoff for inclusion in the final analysis was set as genes with a value of absolute log_2_ fold-change (log_2_FC) ≥ 0.585 and adjusted *p*-value ≤ 0.05. Adusted p-values designated as 1.0 in the raw data indicate the gene could not be included or excluded as significant (generally due to insufficient reads in either the control or experimental samples).

## ACKNOWLEDGEMENTS

We thank Veronica Chemelewski for critical reading of the manuscript, Morgan Chapman for generating the P_*hy*_-*uppS* strain, Tatiana Castro Padovani for assistance in visually screening deletion mutants for loss of Mg_2+_-responsiveness, and Dr. Leendert Hamoen for sharing the Gintool starter package. This work was supported by T. Guo teaching every semester and scrap funds from the startup account of J.K. Herman. We dedicate this work to Ry Young in honor of his retirement.

## Supplementary materials

**Text S1**. Descriptions of strain construction

**Figure S1.**
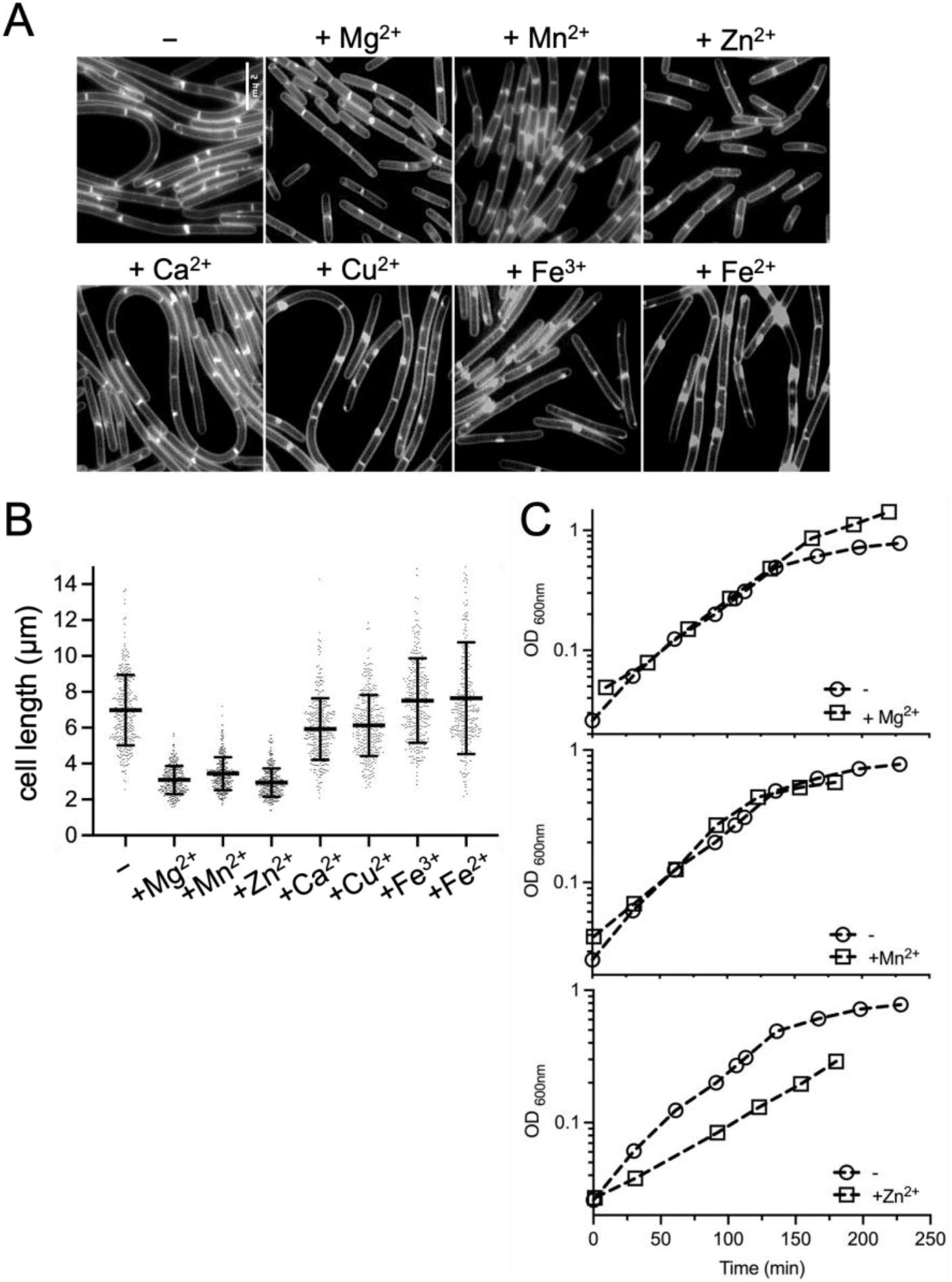
Effect of divalent metals on cell length. Wild-type 168 (BJH004) cells were cultured in CH* without or with the following concentrations of divalent cation salts, as indicated: 10 mM MgCl_2_, 0.1 mM MnSO_4_, 0.2 mM ZnCl_2_, 1.0 mM CaCl_2_, 0.5 mM CuSO_4_, 0.1 mM Fe_2_(SO_4_)_3_, and 0.5 mM FeSO_4_. (A) Micrographs following membrane staining and epifluorescent microscopy (scaled identically). (B) Scatter plots showing the distribution of cell lengths quantitated for 300 cells from each condition. Bars represent the means of 300 cells ± SD. (C) Representative growth curves.

**Figure S2.**
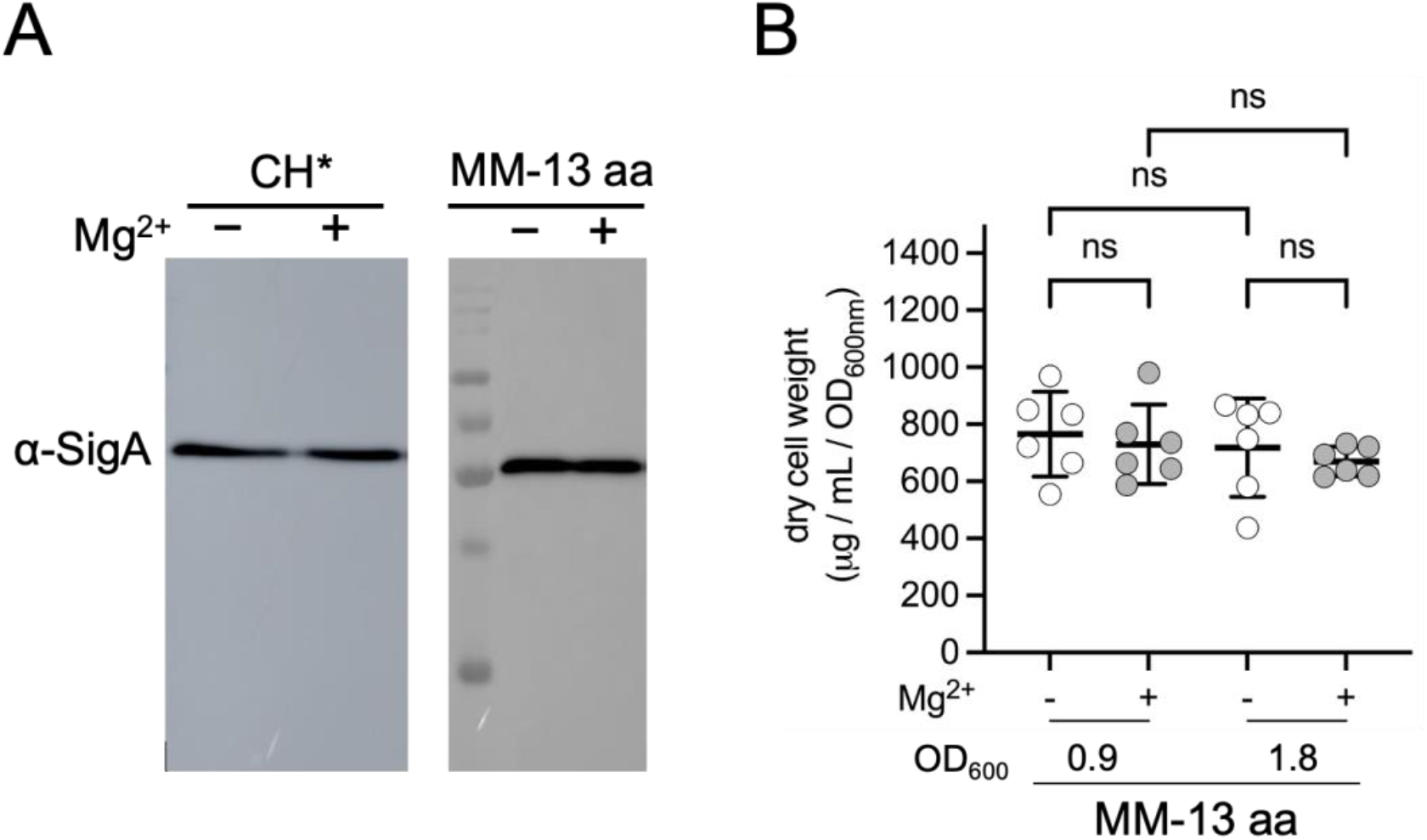
Controls assessing the robustness OD_600_ reading as a measurement of growth. (A) Immunoblot analysis of SigA expression level of *B. subtilis* 168 (BJH004) cells grown in CH* or MM-13 aa. 10.0 mM MgCl_2_ was added as indicated. Cells were harvested at OD_600_ of ∼0.2 and sample loads were normalized to each other using OD_600_ values. (B) Dry cell weight of *B. subtilis* 168 (BJH004) cultured in MM-13 aa. 10.0 mM MgCl_2_ was added as indicated. Growth is still linear at an OD_600_ of 0.9 (see Fig 3A). Statistical analysis was done with Ordinary one-way ANOVA. (ns, p>0.05).

**Figure S3.**
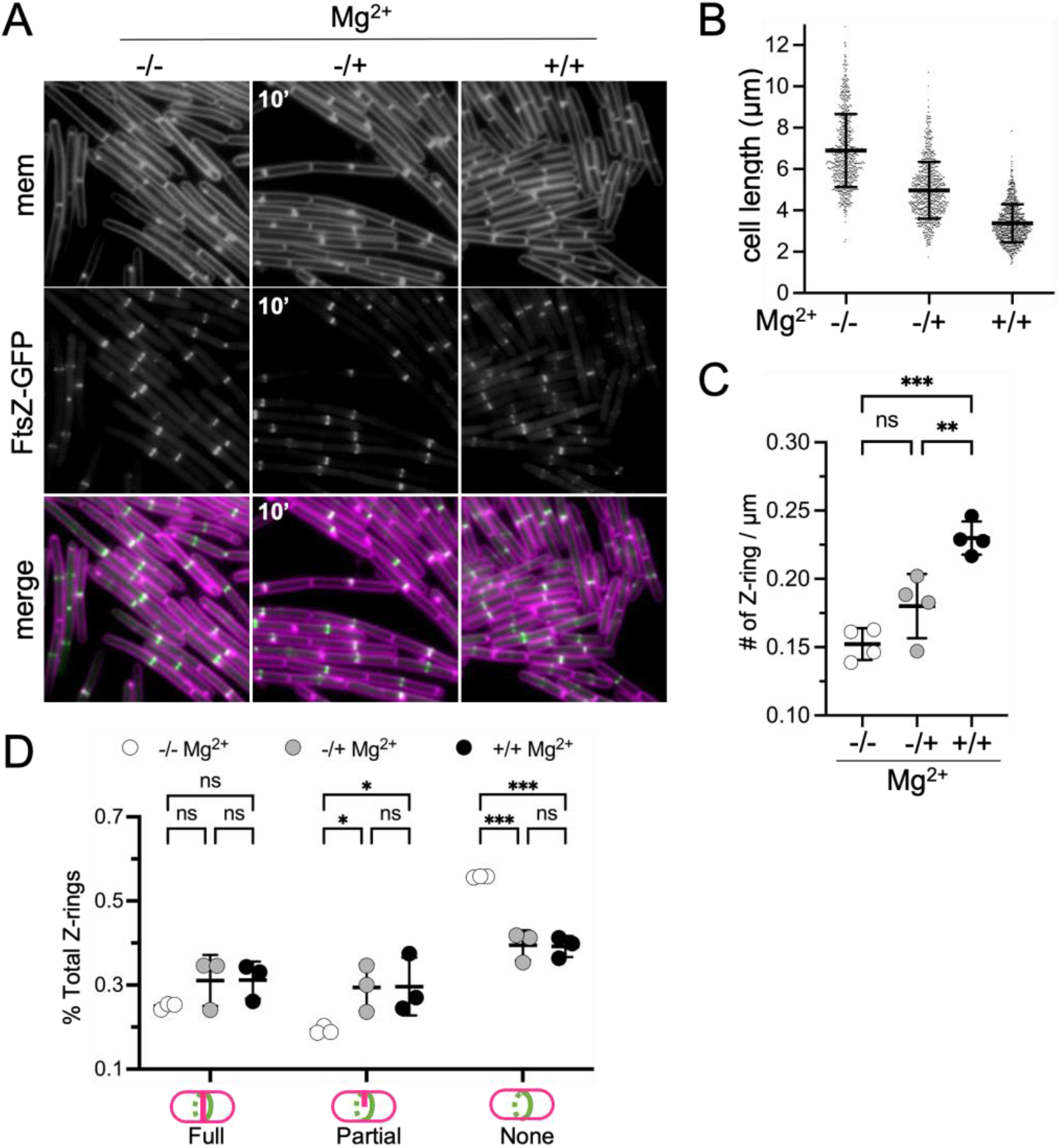
Imaging and quantitation of FtsZ-GFP following Mg_2+_ supplementation. *P*_*spac*_*-ftsZ-GFP* (BTG185) was grown in MM-13aa with 5.0 µM IPTG. For the transfer experiment, cells were imaged 10 min after the addition of 10.0 mM MgCl_2_. (A) Representative micrographs. Membranes stained with FM4-64. (B) Scatter plots showing the distribution of cell lengths. Bars represent the mean cell lengths 800 cells ± SD. (****) p<0.0001. (C) Average number of Z-rings per unit cell length ± SD. Each circle represents the mean of 200 cells from four independent biological replicates per condition. (D) Average fraction of cells with coalesced FtsZ-GFP presenting the indicated septum type ± SD. Each circle represents the mean of 500 cells from three independent biological replicates per condition. (*** 0.0001<p<=0.001), (*, 0.01<p<=0.05), (ns, p>0.05).

**Figure S4.**
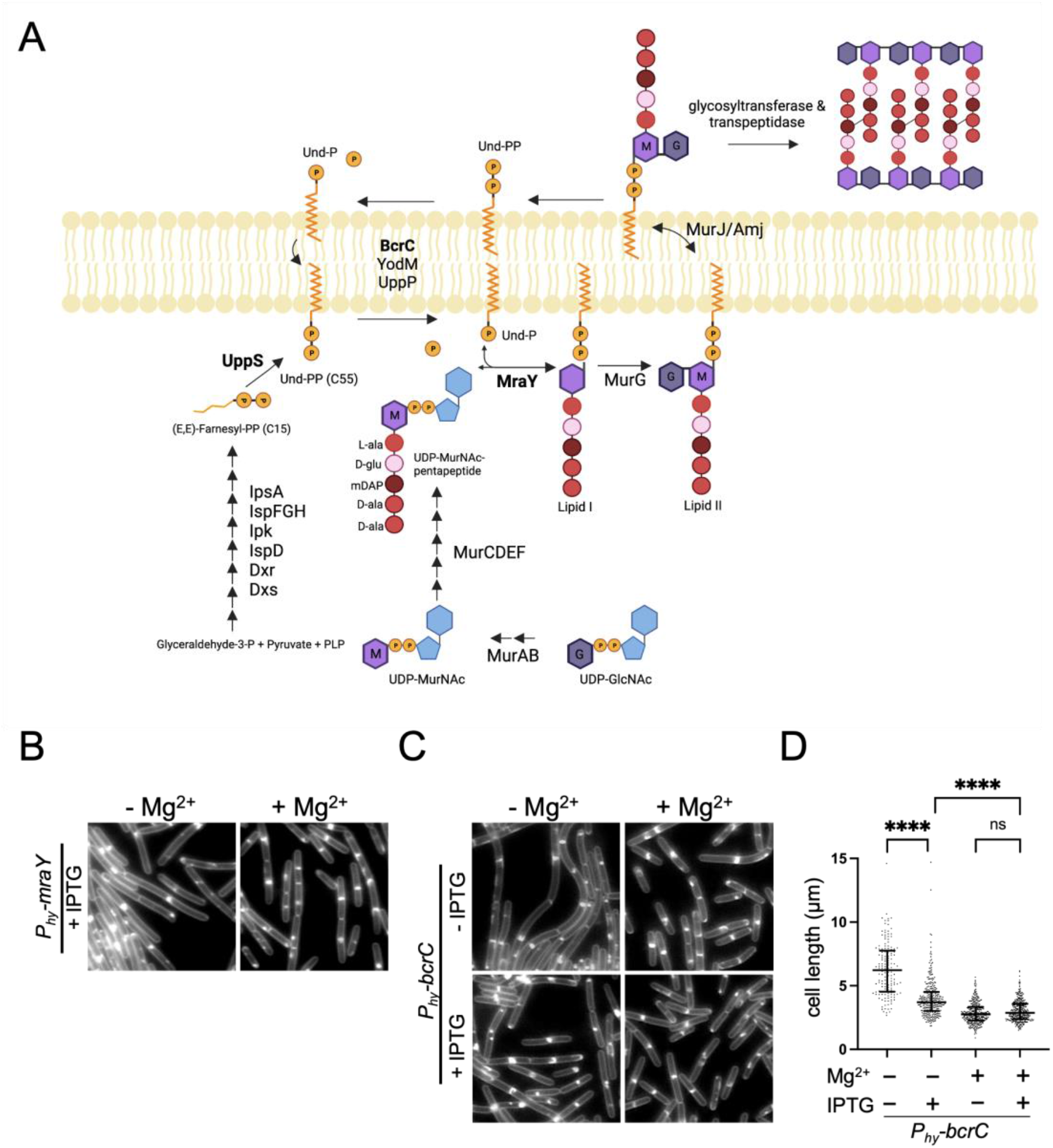
(A) Peptidoglycan biosynthesis pathway. Representative micrographs of (B) *P*_*hy*_*-mraY* (BTG697) and (C) *P*_*hy*_*-bcrC* (BTG678) cultured in CH* with 1.0 mM IPTG and with or without 10.0 mM MgCl_2_. Membranes are stained with TMA. Images are scaled identically. (D) Scatter plots showing the distribution of cell lengths quantitated for 150 cells from each condition. Bars represent the mean cell length ± SD (Ordinary one-way ANOVA). (****)p<0.0001, (ns)p>0.01234. (C) Representative growth curves.

**Table S1.**
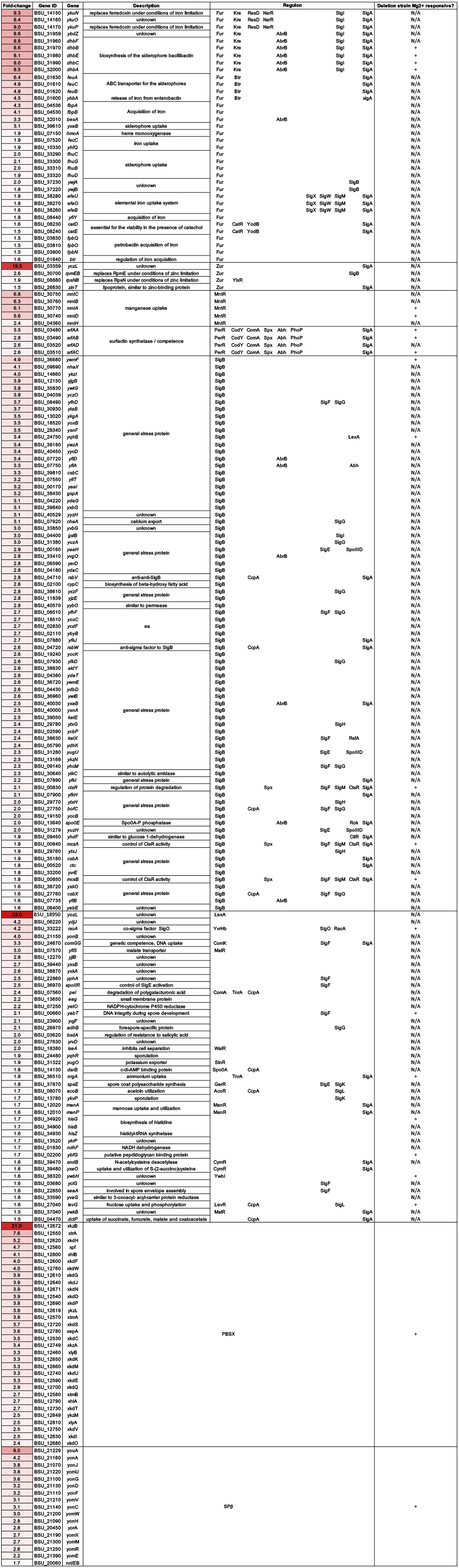
RNA-seq analysis of genes upregulated in (CH* + 10.0 mM MgCl2)/(CH*)

**Table S2.**
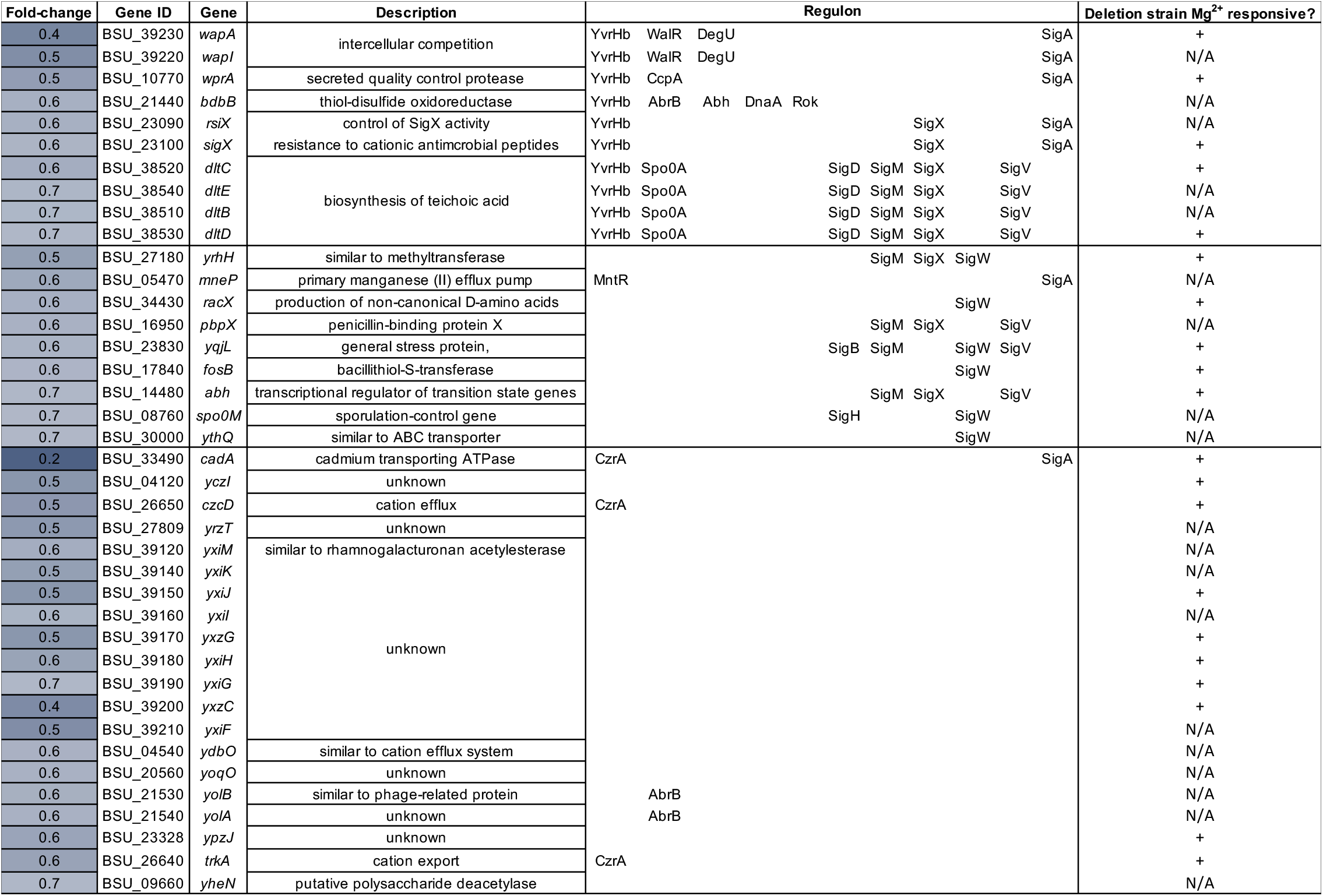
RNA-seq analysis of genes downregulated in (CH* + 10.0 mM MgCl2)/(CH*).

**Table S3.** Deletion strains that retained the Mg^2+^-dependent cell shortening phenotype

|  |  |  |  |  |  |
| --- | --- | --- | --- | --- | --- |
| strain* | genotype | BKK strain | function | Description | Deletion strain Mg<sup>2+</sup> responsive? |
| BJH582 | $\Delta dctS$ | BKK04450 | kinase | regulation of the *dctS*-*dctR*-*dctP* operon | + |
| BJH583 | $\Delta glnK$ | BKK02440 | kinase | regulation of the *glsA*-*glnT* operon | + |
| BJH585 | $\Delta ydfI$ | BKK05420 | regulator | control of *ydfJ* expression | + |
| BJH586 | $\Delta kinC$ | BKK14490 | kinase | initiation of sporulation | + |
| BJH587 | $\Delta cheA$ | BKK16430 | kinase | modulation of flagellar switch bias | + |
| BJH588 | $\Delta lytT$ | BKK28920 | regulator | control of pyruvate utilization | + |
| BJH589 | $\Delta desK$ | BKK19190 | kinase | regulation of cold shock expression of *des* | + |
| BJH590 | $\Delta ybdK$ | BKK02010 | kinase | unknown | + |
| BJH591 | $\Delta ykoH$ | BKK13260 | kinase | unknown | + |
| BJH592 | $\Delta yhcY$ | BKK09320 | kinase | unknown | + |
| BJH593 | $\Delta bceS$ | BKK30390 | kinase | resistance against toxic peptides | + |
| BJH594 | $\Delta citS$ | BKK07580 | kinase | regulation of citrate uptake | + |
| BJH595 | $\Delta yxjL$ | BKK38910 | regulator | unknown | + |
| BJH596 | $\Delta yesM$ | BKK06950 | kinase | unknown | + |
| BJH597 | $\Delta yrkQ$ | BKK26420 | kinase | unknown | + |
| BJH598 | $\Delta liaS$ | BKK33090 | kinase | regulation of the *liaI*-*liaH*-*liaG*-*liaF*-*liaS*-*liaR* operon | + |
| BJH599 | $\Delta lnrJ$ | BKK08290 | kinase | resistance to linearmycin | + |
| BJH600 | $\Delta natK$ | BKK02730 | kinase | regulation of the *natA*-*natB* operon | + |
| BJH601 | $\Delta yclK$ | BKK03760 | kinase | unknown | + |
| BJH602 | $\Delta resE$ | BKK23110 | kinase | regulation of aerobic and anaerobic respiration | + |
| BJH605 | $\Delta yxjM$ | BKK38900 | kinase | unknown | + |
| BJH609 | $\Delta comP$ | BKK31690 | kinase | regulation of genetic competence and quorum sensing | + |
| BJH610 | $\Delta spo0A$ | BKK24220 | regulator | initiation of sporulation | + |
| BJH611 | $\Delta cssS$ | BKK33020 | kinase | control of cellular responses to protein secretion stress | + |
| BJH612 | $\Delta degS$ | BKK35500 | kinase | regulation of degradative enzymes, genetic competence, and other adaptive responses | + |
| BJH613 | $\Delta yvrG$ | BKK33210 | kinase | regulation cell surface maintenance | + |
| BJH615 | $\Delta resD$ | BKK23120 | regulator | regulation of aerobic and anaerobic respiration | + |
| BJH616 | $\Delta ypsdS$ | BKK34710 | kinase | resistance against toxic peptides | + |
| BJH617 | $\Delta yvfT$ | BKK34070 | kinase | unknown | + |
| BJH618 | $\Delta psdR$ | BKK34720 | regulator | resistance against toxic peptides | + |
| BJH621 | $\Delta glnL$ | BKK02450 | regulator | regulation of the *glsA*-*glnT* operon | + |
| BJH622 | $\Delta cheY$ | BKK16330 | regulator | modulation of flagellar switch bias | + |
| BJH623 | $\Delta yneI$ | BKK17940 | regulator | unknown | + |
| BJH624 | $\Delta lytS$ | BKK28930 | kinase | control of pyruvate utilization | + |
| BJH626 | $\Delta ydfH$ | BKK05410 | kinase | control of *ydfJ* expression | + |
| BJH638 | $\Delta malK$ | BKK31520 | kinase | regulation of malate uptake | + |
| BAS206 | $\Delta 6$ | | | *Bacillus subtilis* 168 $\Delta SP\beta$ $\Delta skin$ $\Delta PBSX$ $\Delta prophage$ 1 pks::cat $\Delta prophage$ 3 | + |

\*All BKK-based strains were generated by introducing genomic DNA from the indicated BKK strain into *B. subtilis* 168 competent cells and selecting for kanamycin resistance.

**Table S4.** Strains.

| Strain | Wildtype (WT) Strains | Reference |
| --- | --- | --- |
| BJH004 | *Bacillus subtilis* 168 *trpC2* | Bacillus Genetic Stock Center 1A1 |
| BJH001 | PY79 | Obtained from David Rudner (1) |
| BJH403 | 3610 | Dan Kearns (2) |
| BTG169 | *Bacillus subtilis* 168 *trpC+* | This study |
| BAS206 | $\Delta 6$ ($\Delta SP\beta$ $\Delta skin$ $\Delta PBSX$ $\Delta prophage$ 1 *pks::cat* $\Delta prophage$ 3) | Obtained from Ryland Young who acquired from Dean Scholl (3, 4) |
| ***Bacillus subtilis* 168 derivatives** |  |  |
| BTG182 | *amyE::P<sub>mgtE</sub> leader (nt -486 - +25)-lacZ (cat)* | Wade Winkler (5) |
| BTG333 | *mpfA::kan, amyE::P<sub>mgtE</sub> leader (nt -486 - +25)-lacZ (cat)* | This study |
| BTG186 | *amyE::P<sub>xyl</sub>-gfp-zapA (cat)* | This study |
| BTG185 | *amyE::P<sub>spac</sub>-ftsZ-gfp (cat)* | This study |
| BTG708 | *amyE::P<sub>hy</sub>-uppS (spec)* | This study |
| BTG697 | *amyE::P<sub>hy</sub>-RBS- mraY (spec)* | This study |
| BTG678 | *amyE::Phy-RBS-bcrC (spec)* | This study |

**Table S5.** Plasmids.

| Plasmid | Description | Reference |
| --- | --- | --- |
| pDR111 | *amyE::P<sub>hy</sub> (spec)(amp)* | David Rudner |

**Table S6.**
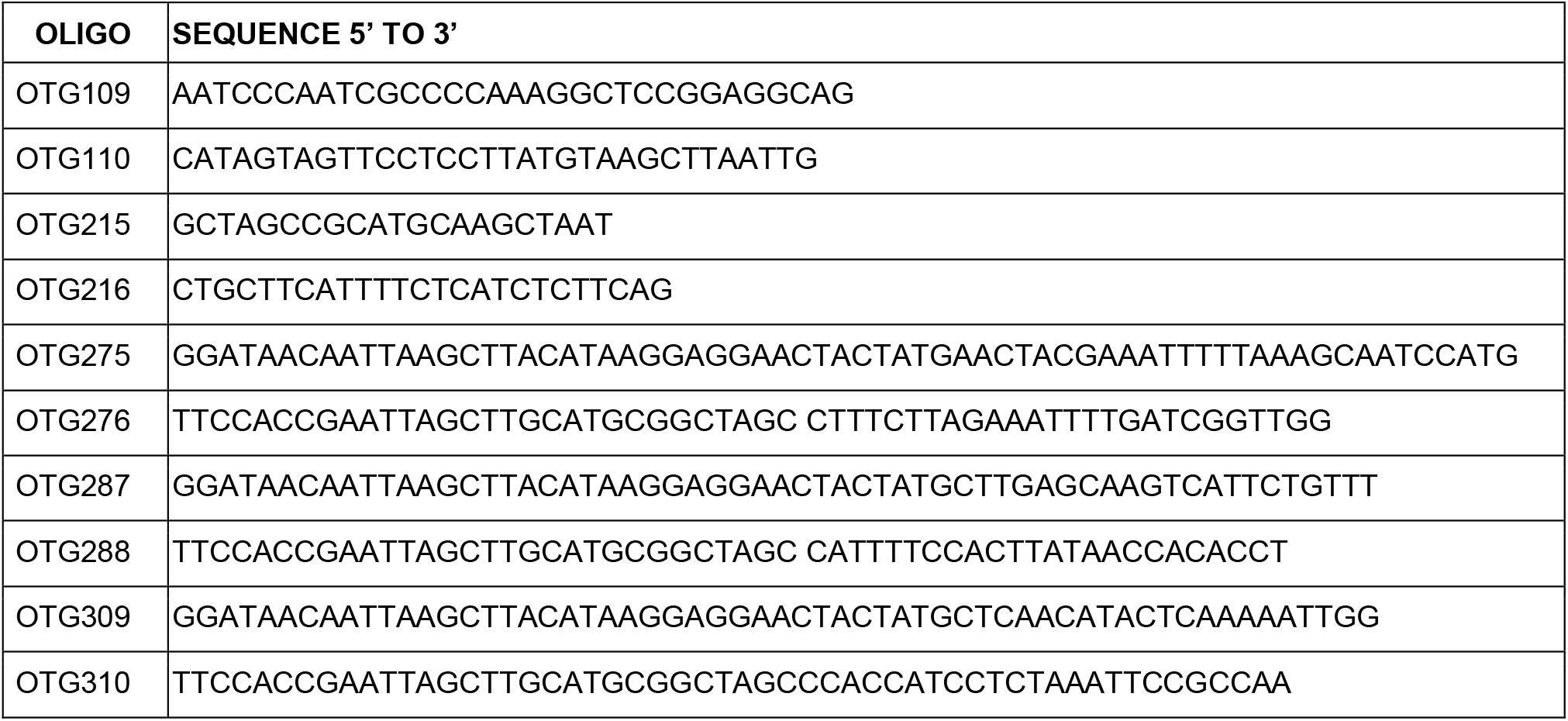
Oligonucleotides.

**Text S1. Strain construction**

**Strain construction** (in alpha-numerical order)

The *B. subtilis* kanamycin resistant deletion strains in Table S3 were derived from the BKK collection (6). Genomic DNA from each BKK strain was transformed into competent BJH004 and selection was performed on LB containing 10 µg ml-1 kanamycin.

**BTG169** *B. subtilis 168 trpC2+* was generated by transforming BJH004 with 40 ng of genomic DNA from PY79 and selecting for growth on minimal medium lacking tryptophan. The strain was confirmed to harbor the PY79 *trpC* allele using whole genome sequencing.

**BTG182** [*amyE::P*_*mgtE*_ *leader (nt -486 - +25)-lacZ (cat)*] was created by transforming *Bs168* with genomic DNA from bCAW2036 [*amyE::P*_*mgtE*_ *leader (nt -486 - +25)-lacZ (cat)*] selecting for growth on LB plates containing 7.5 µg ml-1 chloramphenicol.

**BTG185** [*amyE::P*_*spac*_*-ftsZ-gfp (cat)*] was created by transforming *Bs168* with genomic DNA from gJW28 [*amyE::P*_*spac*_*-ftsZ-gfp (cat)*] selecting for growth on LB plates containing 7.5 µg ml-1 chloramphenicol.

**BTG186** [*amyE::P*_*xyl*_*-gfp-zapA (cat)*] was created by transforming *Bs168* with genomic DNA from gJW29 [*amyE::P*_*xyl*_*-gfp-zapA (cat)*] selecting for growth on LB plates containing 7.5 µg ml-1 chloramphenicol.

**BTG333** [*mpfA::kan, amyE::P*_*mgtE*_ *leader (nt -486 - +25)-lacZ (cat)*] was created by transforming BJH681 [*mpfA::kan*] with genomic DNA from bCAW2036 [*amyE::P*_*mgtE*_ *leader (nt -486 - +25)-lacZ (cat)*] selecting for growth on LB plates containing 7.5 µg ml-1 chloramphenicol.

**BTG678** [*amyE::P*_*hy*_*-bcrC (spec)*] was created by transforming *Bs168* with a linear Gibson assembly product encoding three fragments including a region upstream of *amyE* with a spectinomycin cassette using oTG109 and oTG110, the *P*_*hyperspank*_*-bcrC* using oTG275 and oTG276, a region downstream of *amyE* into *B. subtilis* 168 using oTG215 and oTG216 and selecting for for growth on LB plates containing 100 µg ml-1 spectinomycin. The detailed protocol is available in reference (7).

**BTG697** [*amyE::P*_*hy*_*-RBS-mraY (spec)*] was created by transforming *Bs168* with a linear Gibson assembly product encoding three fragments including a region upstream of *amyE* with a spectinomycin cassette using oTG109 and oTG110, the *P*_*hyperspank*_*-mraY* using oTG287 and oTG288, a region downstream of *amyE* into *B. subtilis* 168 using oTG215 and oTG216 and selecting for for growth on LB plates containing 100 µg ml-1 spectinomycin. The detailed protocol is available in reference (7).

**BTG708** [*amyE::P*_*hy*_*-uppS (spec)*] was created by transforming *Bs168* with a linear Gibson assembly product encoding three fragments including a region upstream of *amyE* with a spectinomycin cassette using oTG109 and oTG110, the *P*_*hyperspank*_*-uppS* using oJH309 and oJH310, a region downstream of *amyE* into *B. subtilis* 168 using oTG215 and oTG216 and selecting for for growth on LB plates containing 100 µg ml-1 spectinomycin. The detailed protocol is available in reference (7).

